# Parallel factor analysis enables quantification and identification of highly-convolved data independent-acquired protein spectra

**DOI:** 10.1101/2020.04.21.052654

**Authors:** Filip Buric, Jan Zrimec, Aleksej Zelezniak

**Affiliations:** Department of Biology and Biological Engineering, Chalmers University of Technology, Kemivägen 10, SE-412 96, Gothenburg, Sweden; Science for Life Laboratory, Tomtebodavägen 23a, SE-171 65, Stockholm, Sweden

**Keywords:** mass spectrometry, proteomics, deconvolution, PARAFAC, tensor algebra

## Abstract

High-throughput data-independent acquisition (DIA) is the method of choice for quantitative proteomics, combining the best practices of targeted and shotgun proteomics approaches. The resultant DIA spectra are, however, highly convolved and with no direct precursor-fragment correspondence, complicating the analysis of biological samples. Here we present PARADIAS (PARAllel factor analysis of Data Independent Acquired Spectra), a GPU-powered unsupervised multiway factor analysis framework that deconvolves multispectral scans to individual analyte spectra, chromatographic profiles, and sample abundances, using the PARAFAC tensor decomposition method based on variation of informative spectral features. The deconvolved spectra can be annotated with traditional database search engines or used as a high-quality input for *de novo* sequencing methods. We demonstrate that spectral libraries generated with PARADIAS substantially reduce the false discovery rate underlying the validation of spectral quantification. PARADIAS covers up to 33 times more total ion current than library-based approaches, which typically use less than 5 % of total recorded ions, thus allowing the quantification and identification of signals from unexplored DIA spectra.

## Introduction

The ideal proteomic method should precisely quantify large sets of proteins across multiple samples. To this end, data-independent acquisition (Venable *et al*, 2004) (DIA) is an effective compromise between targeted proteomics using selected reaction monitoring (SRM) and label-free shotgun proteomics with data-dependent acquisition (DDA), combining the respective benefits of high accuracy and consistency (Vowinckel *et al*, 2018; Rosenberger *et al*, 2017) with high-throughput (Messner *et al*, 2019). Multiple issues are addressed, such as the inconsistent quantification due to stochasticity between runs, noticeable especially in DDA experiments with large sample series (Collins *et al*, 2017; Zhang *et al*, 2016). Despite this, an inherent issue is related to the exhaustive fragmentation of the specific mass range using defined isolation windows or “swaths” (Gillet *et al*, 2012). Due to the width of these windows, fragment signals are highly overlapped or “convolved”, with multiple precursors falling in the same window, producing a set of highly overlapping ion mass spectra (Pappireddi *et al*, 2019; Peckner *et al*, 2018; Demichev *et al*, 2019). A computational solution to deconvolve such data would expand the coverage and efficacy of the DIA approach. Thus, development of novel data analysis approaches is currently among major goals in high-throughput proteomics.

The current standard approach for DIA analysis is targeted quantification of the acquired fragment data using spectral libraries containing fragmentation information for a particular peptide (Bruderer *et al*, 2015; Demichev *et al*, 2019; Röst *et al*, 2014; Peckner *et al*, 2018). Library generation is however time-consuming, specific to the instrument, chromatography, and experimental condition, ideally requiring physical sample fractionation complemented with shotgun spectra acquisition (Schubert *et al*, 2015). Another limitation is that only a small portion of analytes are recovered, especially when library generation is based on data-dependent acquisition of relatively few selected high intensity precursors (Ludwig *et al*, 2018; Deutsch *et al*, 2018). Thus, the targeted search for DDA precursor fragments does not take full advantage of resulting digital records of all ions in scans generated in a data-independent manner (Gillet *et al*, 2012). Recent approaches based on large synthetic peptide libraries enable accurate prediction of peptide spectra directly from sequence data (Gessulat *et al*, 2019; Gabriels *et al*, 2019). Computational approaches that utilize MS1 - MS2 co-elution information to generate pseudo-spectra do not require creating experimental libraries (Tsou *et al*, 2015; Wang *et al*, 2015; Li *et al*, 2015). These, however, suffer from the same overlapping fragment signal problem inherent to DIA, which is addressed using heuristics such as interference correction (Bao *et al*, 2013; Keller *et al*, 2015; Demichev *et al*, 2019).

Multiway tensor decomposition and other so-called “matrix methods” (Likić, 2009), such as Parallel Factor Analysis (PARAFAC) (Harshman & Others, 1970; Carroll *et al*, 1970; Bro & Others, 1997), use the entire acquired data to extract individual analyte signals and have have been used for over four decades in mass spectrometry and other analytical technologies (Likić, 2009). PARAFAC enables decomposition of multiway data arrays and facilitates the identification and quantification of independent underlying signals, termed “components”, from convolved spectral data. Conveniently, DIA data can be naturally represented as a three-dimensional array or tensor, resulting from the linear combination of individual peptide mass spectra, their elution profiles, and their relative sample contribution, making it amenable to PARAFAC decomposition. However, given the sheer size of DIA proteomics datasets, where an experiment of 100 samples can easily generate more than half a terabyte of numerical data, computational decomposition of DIA proteomics data using conventional CPU-based multiway analysis frameworks (Andersson & Bro, 2000) is not feasible. Furthermore, existing PARAFAC applications usually involve smaller data sets consisting of at most a few hundred known analytes, so far limiting PARAFAC applications to relatively simple computational problems. On the other hand, with DIA proteomics data, one deals with an unknown set of tens of thousands of analytes, thus requiring a way to search a much larger model space than is currently achievable.

Hence, here we present a GPU-accelerated multiway tensor decomposition approach called *PARAllel factor analysis of Data-Independent Acquired Spectra* (PARADIAS), consisting of a data decomposition pipeline that enables spectra retrieval and quantification of analytes directly from DIA data. By using a data partitioning scheme and relying on the massive parallelism of modern GPUs, we achieve a technical leap, enabling untargeted decomposition of very large, high-throughput proteomics data. Moreover, the central method to the pipeline, PARAFAC, does not require *a priori* spectral information about the analytes in order to perform the decomposition. The individual deconvolved spectra produced by our pipeline may be analyzed with conventional peptide search engines (Park *et al*, 2008; Deutsch *et al*, 2015; Kim & Pevzner, 2014) to produce peptide-spectrum matches (PSMs) for building high-accuracy spectral libraries. We show that, by using PARADIAS, we can extract up to 33 times more analyte signal from DIA scans compared to library-based approaches, and, moreover, cover the entire m/z space of scans, enabling the usage of the majority of noise-accounted signal ions obtained from a sample. We also demonstrated that spectra recovered by PARADIAS circumvent the problem of false quantifications, a major challenge present in targeted DIA proteomics.

## Results

### PARADIAS: A GPU-accelerated software pipeline for deconvolving DIA data

We developed the PARADIAS pipeline, capable of recovering spectral features in unsupervised fashion and computationally feasible time (Figure 1a), by leveraging the power of the modern tensor algebra frameworks PyTorch (Paszke *et al*, 2017) and Tensorflow (Abadi *et al*, 2016), which take advantage of the parallelism and throughput of floating point operations in graphic processing units (GPU), as well as the distributed Big Data computing framework Apache Spark (Zaharia *et al*, 2016). Briefly, PARADIAS partitions all provided DIA scans into a collection of small, independent tensors, corresponding to precursor isolation windows and time intervals (Methods M1). It then performs multiple decompositions of each of these tensors in parallel, accounting for a range of possible numbers of components in each tensor (Figure 1a, Methods M2). As the observed intensities in DIA LC-MS/MS scans result from linear combinations of individual fragmented peptide spectra, their elution profiles, and their relative abundance across all samples, each PARAFAC component ideally represents an analyte as a triplet of its m/z spectrum, retention time peak and relative sample contribution (Figure 1b). The decomposition results are then refined by selecting the best models based on the quality of reconstructed signals, i.e. the unimodality of the elution profile. (Methods M3, Supplementary Note S1-1). A critical step in constructing a PARAFAC model is deciding *a priori* the number of components F, complicated by the fact that PARAFAC models do not “nest”, i.e. a model for F+1 is not simply a model for F with an extra component (Smilde *et al*, 2005). Deciding the value automatically is generally an open problem (Liu *et al*, 2016), and the various diagnostics and procedures used to this end (Bro & Kiers, 2003) often require human verification, which is not feasible for data-rich proteomics workflows, with hundreds of thousands of models that need to be examined. Our approach therefore exhaustively constructs all possible models within the expected range, and then uses the shape of the resulting elution profiles and accounts for noise to automatically select valid models, resulting in optimal precursor identifications. The recovered m/z spectra (Figure 1c) can be directly searched using standard tool sets such as Crux (McIlwain *et al*, 2014), TPP (Deutsch *et al*, 2015), and MS-GF+ (Kim & Pevzner, 2014) to (i) produce peptide-spectrum matches (PSMs) (Figure 1d), (ii) build spectral libraries (Methods M4), (iii) be used for *de novo* sequencing, or (iv) be used directly as linearly independent features for machine learning applications.

**Figure 1.**
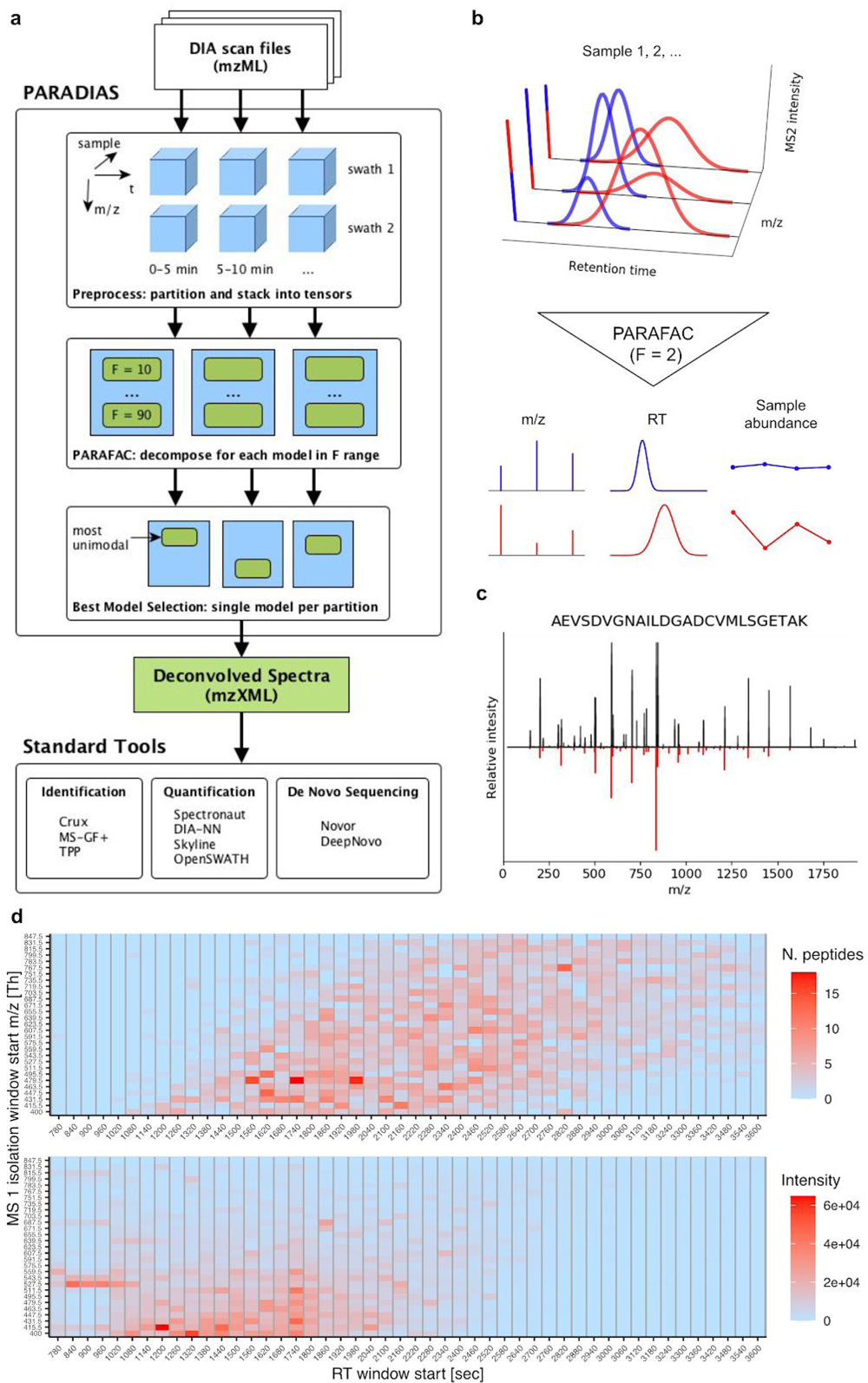
The PARADIAS pipeline, illustration of PARAFAC decomposition, and example results. (a) High-level structure of the PARADIAS framework. Under the hood, PARADIAS uses tools typically applied for processing of big data (on the order of hundreds of GB of numerical data) to perform PARAFAC decomposition of similarly large DIA data. It operates in a parallelized way, employing tensor computation frameworks that leverage the speed of GPU cards for these types of matrix operations. PARADIAS takes in all provided DIA scan files together, partitions them into a collection of independent tensors according to swath and retention time windows, then performs multiple decompositions of each of these tensors, accounting for a range of possible number of components, to account for an unknown number of peptides in each partition. The best models are selected such that most components have unimodal elution profiles (Methods M3, Figure 3). PARADIAS output consists of a file in mzXML format containing the deconvolved spectra. This file is orders of magnitude smaller than the input scan files, which speeds up downstream analytical methods. (b) Conceptual illustration of the PARAFAC decomposition method for two components. Acquired DIA signals can be expressed as a linear combination of individual peptide mass spectra, their elution profiles, and their relative sample contribution. PARAFAC considers all sample scans at once and decomposes the three-dimensional (m/z, retention time, sample) tensor structure into deconvolved components. (c) Example of PARAFAC output spectrum matched to a peptide by Comet. Theoretical spectrum (predicted with Prosit (Gessulat *et al*, 2019)) of the peptide (black) is plotted against fragments matched (66%) to the deconvolved spectrum output by the pipeline (red). (d) Peptide identification using Crux and MS-GF+ on PARADIAS output (top) largely matches the distribution of input DIA scan MS 1 intensities (bottom, single yeast lysate scan). Peptide count and intensities are shown per retention time (RT) and precursor isolation windows, according to the pipeline’s data partitioning scheme. RT windows are highlighted with light gray vertical lines. The horizontal streaks that show up in the ranges 11 - 17 minutes and 527 - 559 m/z (lower-left) are likely contaminants (e.g. nothing was identified by Spectronaut (FDR < 5%) in this range) and are not reflected in any PARADIAS identifications.

### Precise protein identification and quantification with PARADIAS

We first evaluated whether PARADIAS-deconvolved spectra were identifiable by conventional peptide search engines. First, we tested our approach on a *S. cerevisiae* lysate dataset (Vowinckel *et al*, 2018), referred to here as *yeast replicates*, which consisted of 9 consecutive injections acquired in SWATH mode on a conventional Sciex 5600 QqTOF instrument with microflow setup. To identify the precursors, PARADIAS solved a total of 176175 PARAFAC models (29 swaths x 75 1-min windows x 81 models) on a standard GPU-equipped workstation. As a benchmark for comparison, we used DIA-Umpire (Tsou *et al*, 2015), a widely used tool for building spectral libraries directly from DIA data, and considered results from Crux (Comet coupled with Percolator) and MS-GF+ search engines separately, to assess their performance. Whilst MS-GF+ and Crux respectively identified a total of 1032 and 2014 proteins using the DIA-Umpire pseudo-spectra produced for each yeast replicate, in contrast, only 489 and 684 proteins (1750 and 1553 peptides), respectively, were identified using the same search tools on the output from PARADIAS. However, when considering only the proteins that appear in at least 8 of the technical replicates, as low as 66 and 583 proteins were found at 1% FDR using DIA-Umpire in conjunction with MS-GF+ and Crux respectively, pointing at false positive identifications (Figure 2a). PARAFAC decomposition, in contrast, captures the same analytes across all input samples, thus the commonality of high-confidence protein IDs across technical replicates is inherent to the method. Overall, 53% overlap, consisting of 439 common proteins, was detected between PARADIAS and DIA-Umpire coupled with Crux (considering only proteins identified in at least 8 samples), with 245 proteins unique to the former (Figure 2a). The peptide quantifications (Figure 2b) based on the reconstructed spectral library are precise (median CV = 9.3% for the yeast replicates dataset) and reproducible (mean Pearson’s *r* = 0.99 and *p*-value < 1e-16 between the replicates).

**Figure 2.**
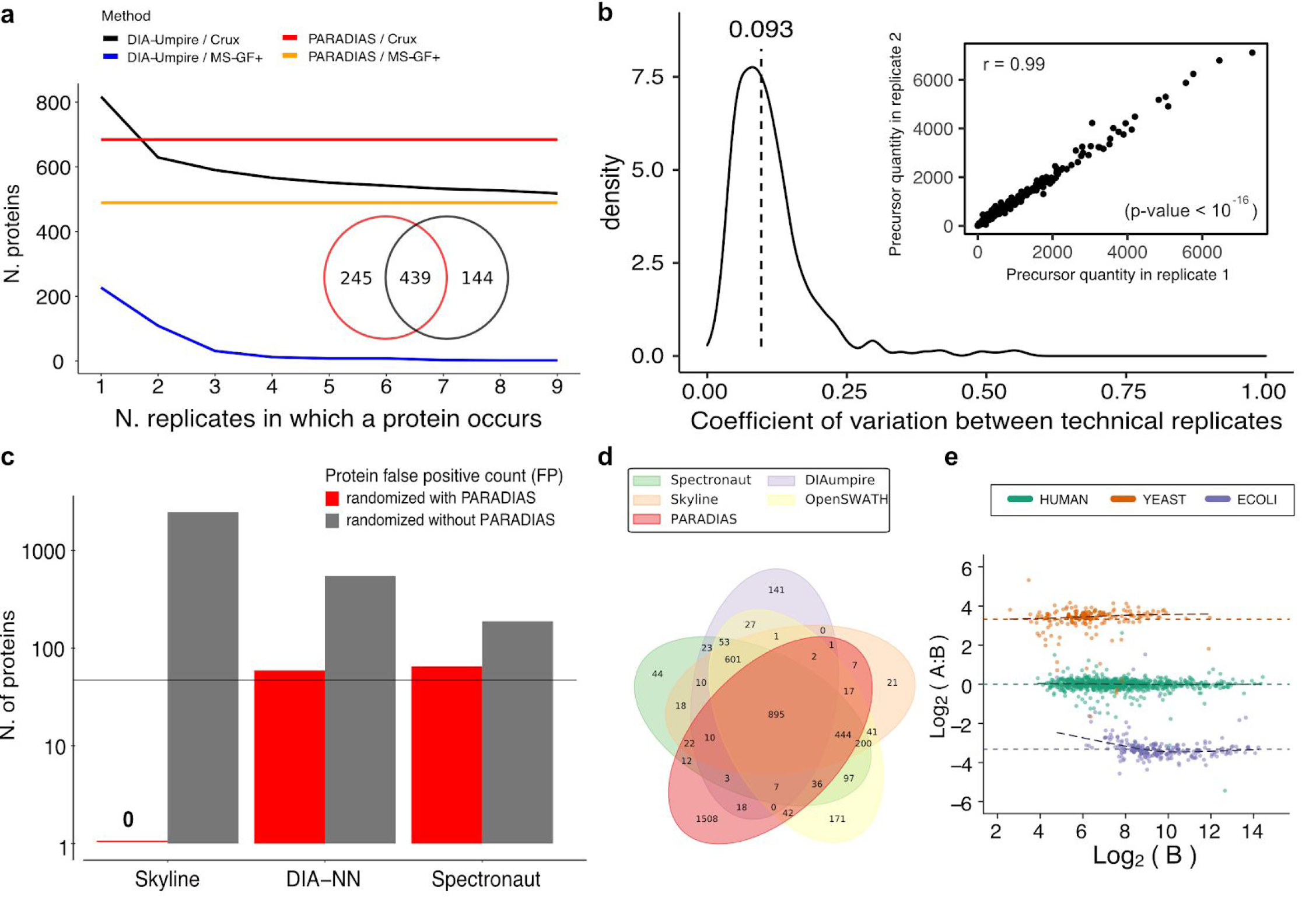
Precise protein identification and quantification with PARADIAS. **(a)** Proteins identified with Crux and MS-GF+ run on DIA-Umpire pseudo-spectra for each replicate, counted according to their prevalence across the replicates, compared with Crux results on PARADIAS output. Strikingly, MS-GF+ identifies very few proteins in all 9 replicates. PARADIAS produces deconvolved spectra from all input replicates, thus the number of IDs reflects the entire dataset. Inset: Overlap of proteins identified by Crux with DIA-Umpire in at least 8 replicates, and proteins identified with PARADIAS. **(b)** Precursor quantity coefficient of variation (median CV = 9.3%, plotted as dashed vertical line) across the yeast replicates dataset, obtained from DIA-NN using a PARADIAS library. Inset: an example of highly correlated quantities between two replicates. **(c)** False positives count underestimation on the yeast replicates dataset, across different software, at 5% FDR. The bar plots show the number of protein IDs under a randomized trial (where all results are false positives), both with and without using PARADIAS as a source of the spectral library, compared to the average expected number of false positives (47) based on baseline (nonrandomized) runs for each tool. **(d)** Overlap between proteins identified with PARADIAS coupled with Crux and MS-GF+ on the LFQbench HYE110 dataset, and published results from other tools. PARADIAS results have 10-fold more unique identifications. **(e)** LFQbench HYE110 results for PARADIAS coupled with DIA-NN, showing quantification of human (green), *S. cerevisiae* (orange), and *E.coli* (purple) peptides. The DIA data are acquired from two hybrid proteome mixtures A and B with known organism concentrations. Plotted are log-transformed ratios (log_2_(A/B)) of peptide concentrations over the log-transformed intensity of sample B, against the expected values for each organism (black dashed lines). Regression curves are shown with black dashed lines.

The inconsistencies in peptide quantifications are often attributed to data-dependent acquisition approaches (Zhang *et al*, 2016), due to their stochastic nature of peptide selection, which is dependent on instrument performance. Although DIA methods typically produce far more complete data matrices (Collins *et al*, 2017), the consistency of identification may be highly dependent on the inference correction procedures and the way false discovery rate is estimated from DIA data. To exemplify this further, we built a library based on *in silico* digestion of a yeast proteome, where we randomly shuffled 50% of amino acids in each peptide sequence. Despite the fact that only peptides which did not exist in the original organism were considered, we quantified 2457, 546, and 188 proteins, using the conventional library-based search tools Skyline (Pino *et al*, 2017), DIA-NN (Demichev *et al*, 2019), and Spectronaut (Bruderer *et al*, 2015), respectively, with 5% FDR (Methods M5). This is on average over 22 times above the expected number of false discoveries (Figure 2c). All of these identifications were attributed solely to the unique fragments arising from the mutated precursors, that would otherwise not be present in the original yeast spectral library. In contrast, quantification with these tools using a library constructed from PARADIAS output spectra yielded an average of 41 identifications, which is below the average expected number of 47 false positives at 5% FDR across all three tools, pointing to the validity of identifications when using PARAFAC-recovered spectra.

We next evaluated whether PARADIAS could resolve spectra in a complex background such as the LFQbench HYE110 dataset (Navarro *et al*, 2016), containing two mixtures with different ratios of human, *Escherichia coli*, and *S. cerevisiae*, the latter two of which are present in quantities close to the limit of detection (5%) alternatively in the two mixtures. As, typically, the number of detected and quantified proteins is highly dependent on the particular tool and FDR estimation method, one would expect to find differences between available tools (Navarro *et al*, 2016) and PARADIAS, given that the latter works by searching against deconvolved spectra rather than matching against scans using a library. Nevertheless, running Crux and MS-GF+ with an FDR threshold of 1% on PARADIAS output spectra yielded a total of 3024 proteins, comparable to the other methods in the benchmark study (Navarro *et al*, 2016), albeit at an overall lower rate of peptide identification (5908 peptides, with a median ratio of peptides to protein of 4, see Figure S1-1). Out of these proteins, 1508 were unique to PARADIAS (Figure 2d), which is 16 higher than the median number of unique proteins of the other methods. We noticed that 601 proteins not identified with PARADIAS were instead found using all methods that, in essence, use the same target-decoy fragment mass search algorithm for FDR estimation (Reiter *et al*, 2011). Analogously to the yeast example above, we built an *in silico* library of randomized peptide sequences, where this time we reduced the amino acid shuffling to 30%, which resulted in a library with no peptides present in the non-randomized protein sequences of any of the three target organisms. The results were different between the tools, with up to 1857 false protein identities found (Figure S1-2). As all false identities were based on fragments that did not originate from the nonrandomized spectral library, considering the false positive rate underestimation of the conventional tools, up to 70% (421 of the 601 common proteins shown in Figure 2d) of proteins identified at 1% FDR with tools other than PARADIAS are thus put into the question. The quantification based on a PARADIAS library, built using conventional methods (Navarro *et al*, 2016; Demichev *et al*, 2019) with an FDR threshold of 1%, resulted in precise ratios between the two mixtures A and B in the dataset (Figure 2e), showing precise quantification using a spectral library constructed directly from the decomposed spectra.

### DIA spectra are still proteomic “dark matter”

Among the advantages of DIA is that acquired data represent a digital snapshot of all ions obtained from a sample (Gillet *et al*, 2012). While the idea is appealing, current library-based methods retrieve only a minor fraction of analytes present in a sample, leaving the majority of acquired DIA spectra unused. To demonstrate this, we calculated the overlap between m/z values of a DIA scan from the HYE110 dataset (Navarro *et al*, 2016) and the corresponding published library. Allowing for a 5-minute retention time (RT) and a 50 ppm m/z tolerance, the spectral library matched 80.77% of the m/z space of this scan. However, summing over the corresponding signal intensities, the library covered only 2.2% of total ion current (TIC) recorded in the scan (Figure 3a), showing that the majority of matched m/z points comprise baseline signals. Indeed, by filtering out all scan intensities below 1, the covered m/z space drops to 3.46%. Thus, the remaining analyte signals, i.e. unlabeled features, are missed by current conventional tools that rely on targeted quantifications. On the contrary, the PARAFAC decomposition method considers virtually all m/z space (Methods M2) in a DIA scan and discards unsystematic noise by leveraging the variability between scans to produce deconvolved spectra (Smilde *et al*, 2005). Thus, a reconstructed scan obtained by recombining PARAFAC output modes (Methods M6) covered 72% of the same HYE110 scan TIC (Figure 3a), which is 33 times higher than what is covered by the spectral library. Analogous results were obtained from *Saccharomyces cerevisiae* lysate and its corresponding spectral library built using fractionation (Vowinckel *et al*, 2018), where the reconstructed PARAFAC pseudo-scan accounted for 12-fold more TIC than the corresponding library (Figure S1-3).

**Figure 3.**
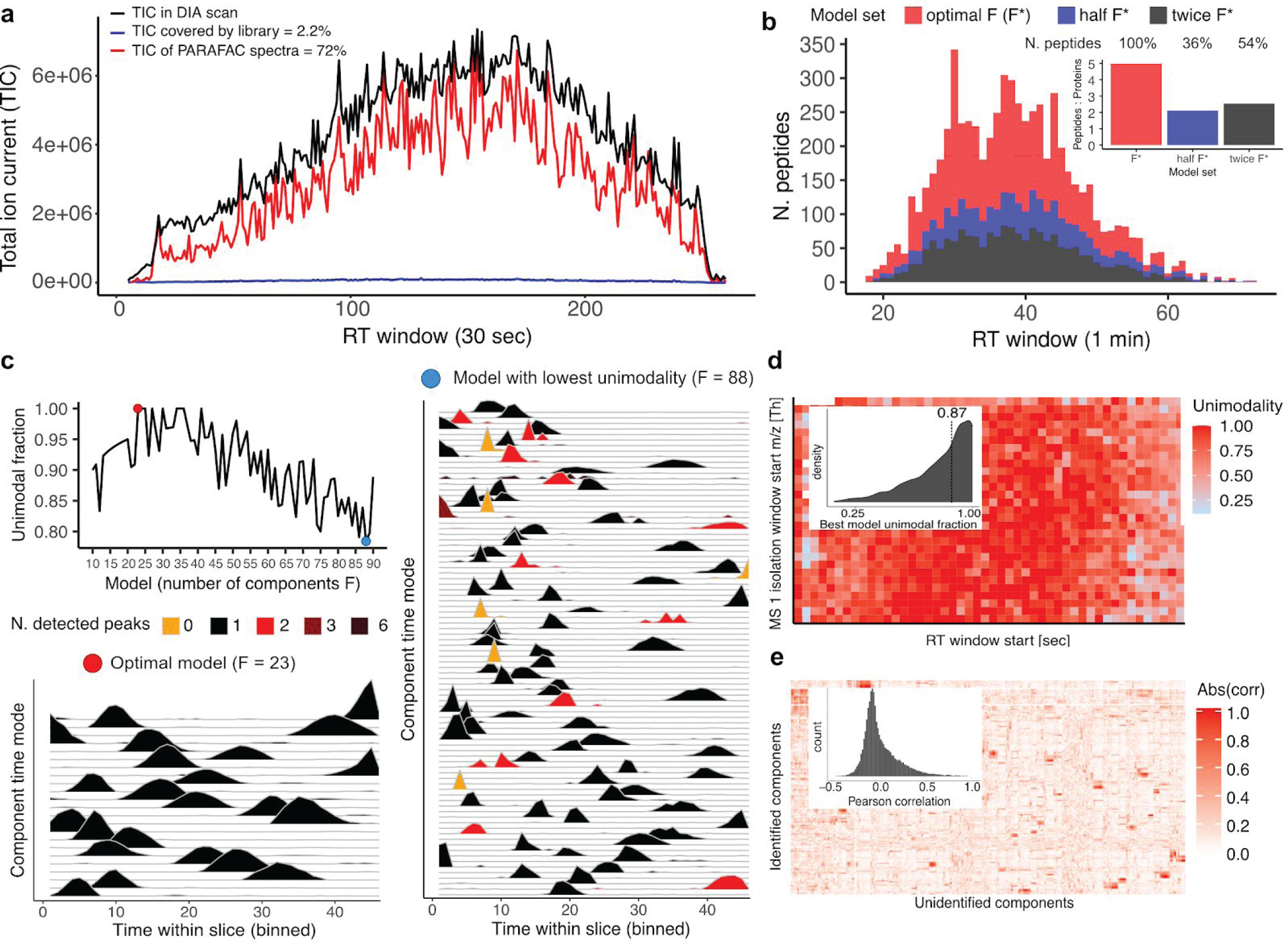
DIA spectra are still proteomic “dark matter”. **(a)** Typical DDA spectral library coverage (2.2%) of the total ion current (TIC) in a centroided DIA scan, compared with that of recovered PARAFAC components (72%). Shown is the TIC per 1 second window of one HYE110 sample scan (black), along with the TIC of the scan spectra matched to the library (blue), and the TIC of a reconstructed scan from PARAFAC output (red), as an approximation of the input DIA scan. Matching with the library allowed for a 5-minute retention time (RT) and a 50 ppm m/z tolerance, and at least 4 library fragments (product m/z points) needed to match for a spectrum to be considered covered. The reconstructed scan was obtained by taking the sum of the outer products of each unimodal component’s m/z and time mode, scaled according to the sample mode coefficient for this scan and the maximum scan intensity (Methods M6). **(b)** Number of peptides identified by Crux in the yeast replicates dataset, using the most unimodal models, compared with two control model sets. The number of Crux (Comet with Percolator) peptide matches at 1% FDR is plotted for models with number components chosen to maximize unimodality fraction (red), and compared against matches for sets of models containing half (blue) and twice (black) the optimal number of components, respectively. In total, 3405 peptides were identified using models chosen based on unimodality, 1222 with models containing half the number of components, and 1837 with models containing twice the number of components. **(c)** Unimodality fractions of all models solved for an example slice of the yeast replicates dataset (MS 1 isolation window 479 - 496 m/z, RT time window 29 - 30 minutes) are shown top-left. The best model (marked with a red dot) was chosen as F = 23, since this is the (lowest) F with 100% unimodal time components. This best model is compared to the worst (F = 88, marked with a blue dot), with the lowest fraction (78%) of unimodal time components. One can see how some of the curves in the worst model are fragmented (higher count of detected peaks) or very narrow (even point-like, resulting in no peak detection). A slice spans 60 seconds and the expected full-width at half maximum (FWHM) of a peak is 12 seconds (Vowinckel *et al*, 2018). We thus considered the non-unimodal components to be noise. **(d)** Heatmap of unimodality fraction across best models for each slice, resulting from the decomposition of the yeast replicates dataset. Inset: Distribution of best model unimodality fraction, with a median of 0.87. **(e)** Heatmap of absolute Pearson correlations between sample modes of identified and unidentified components. Inset: The histogram of Pearson correlations.

Extracting the realistic number of analytes, however, is dependent on the choice of the correct number of components (Smilde *et al*, 2005). The challenge is to identify the number of components that would best match the number of precursors in the samples. Finding the optimal number of components is however an NP-complete problem (Håstad, 1989). Thus, to minimize the risk of overfitting due to an overestimated number of components, we developed an effective empirical approach that functions under the assumption that every analyte has to elute as a single chromatographic peak, our so-called “unimodality criterion” (Methods M3). Indeed, choosing the number of PARAFAC components with most unimodal elution profiles resulted in the highest number of significant (FDR < 1%) precursor identifications (Figure 3b). An increase in number of components did not result in more peptide identifications, rather, it resulted in about half of the amount yielded by the optimal set. This is not surprising, as beyond the correct value, more components will start modeling noise (Smilde *et al*, 2005). As an intuitive example, Figure 3c compares a pair of best and worst models for the yeast replicates dataset, by showing the time modes (elution profiles) of each component in a single slice (single swath and time window), as output by PARAFAC. As demonstrated, the best model captures proper elution curves, whereas non-optimal models comprise a significant number of fragmented or very narrow, single-point curves, representing noise, which is also reflected in 54% lower number of precursor identifications compared to the optimal model (Figure 3b). For the yeast lysate dataset, as the median of the unimodality fraction across best models is 87% (Figure 3d), we attributed the remaining 13% of components to captured noise.

As empirical evidence of the quality of PARADIAS results, the total number of identified proteins for the yeast lysate dataset is comparable to library-based methods (Figures 2a and 2d), and the median number of recovered precursors per protein is 8, which is similar to what is typically expected from these approaches (Navarro *et al*, 2016) (Figure S1-4). Accounting for noisy deconvolutions, the majority (median across datasets = 85%) of recovered spectra (i.e. components) were not mapped to peptide space, and this remaining set of PARAFAC components is overall uncorrelated with the set of identified components (mean Pearson’s *r* = _-_7e-4) (Figure 3e). This was assessed by decomposing the dataset from (Vowinckel *et al*, 2018), consisting of 30 yeast lysate samples from a study of the TOR pathway, and calculating the correlation between identified and unidentified unimodal sample modes (the sample mode of a component holds the contribution of that component to each sample, i.e. analyte abundance in sample). For this dataset, the average unimodal fraction was 90% and, by considering only these non-noisy components, we showed that unidentified components contain non-redundant information that could be leveraged by e.g. machine learning approaches (Zelezniak *et al*, 2018; Haas *et al*, 2017).

Lastly, to extract information from the set of unidentified spectra using existing methods, we searched them for post-translational modifications using MS-GF+ and, moreover, performed *de novo* sequencing using the state of the art machine-learning-based approach DeepNovo (Tran *et al*, 2017), as well as the established tool Novor (Ma, 2015) (Methods M7). *De novo* sequencing benefits from deconvolved input (Muth & Renard, 2018). In total, we identified 186 PTMs in the non-enriched yeast repalicates (Tables S1-1, S1-2), which was twice more than using DIA-Umpire. Moreover, the latter exhibited very low prevalence of identifications, with only 8.2% appearing in at most 2 replicates (Figure S1-6). Likewise, DeepNovo and Novor respectively yielded 4 and 24 times more high confidence (over 80% sequence correctness probability) *de novo* sequences from PARADIAS output, compared to running on DIA-Umpire output (Note S1-2).

## Discussion

Here we presented PARADIAS, a GPU-powered multiway decomposition framework enabling unsupervised and untargeted extraction of analyte signals from DIA data (Figure 1b). PARADIAS solves thousands of decompositions in real time, enabling multiway analyses of dense data-independent-acquired spectra. Parallel factor analysis (Harshman & Others, 1970; Carroll *et al*, 1970), a multiway analysis technique behind PARADIAS, takes advantage of cross-sample analyte variation, enabling deconvolution of mass spectra belonging to individual precursors (Figure 1b, c). The recovered spectra can then be searched using conventional peptide search engines (McIlwain *et al*, 2014; Kim & Pevzner, 2014) or *de novo* sequencing tools (Tran *et al*, 2017; Ma, 2015), to assign analyte identifications. Specifically, we demonstrated PARADIAS quantification precision by building a library from recovered spectra and analysing consecutive injections from yeast lysates acquired using a microflow setup (Vowinckel *et al*, 2018). We also showed PARADIAS performance in quantifying complex background samples, i.e. in an unsupervised fashion, our framework identified peptides and their correct corresponding mixture quantity ratios with high confidence (peptide-level FDR < 1%) in the LFQ benchmark HYE110 dataset (Navarro *et al*, 2016) (Figure 2e). Apart from being a challenging benchmark from an acquisition point of view, as the samples contain a mixture of species in different ratios, correctly identifying peptides and mixture ratios is also not trivial from the data analysis perspective: i) it requires estimating the correct number of components, corresponding to the realistic number of analytes present in the sample; ii) the identification of highly-quality recovered spectra is analogous to data-dependent acquisition directly from MS2 data without MS1 precursor mass isolation; iii) the quantification had to be correct for these recovered spectra in order to yield accurate ratios. Despite these challenges, our unsupervised framework yielded similar results to the established targeted methods (Figure 2).

The quantification and identification of specific analytes requires accurate estimation of false discovery rates, especially crucial when performing unsupervised and untargeted analyte quantification. For DIA data, the procedure is semi-targeted (Ludwig *et al*, 2018), i.e. untargeted acquisition with the targeted analyte quantification either based on an experimental library or *in silico* methods (Yang *et al*, 2020). Conversely, PARADIAS does not depend on a library, but instead builds one using recovered spectra from the observed data, with analyte identification performed *post-hoc* using conventional peptide search engines. Comparison of PARADIAS results to those of other methods showed substantial differences in protein identifications, e.g. in the LFQbench HYE110 dataset, 601 proteins were quantified by all other methods except PARADIAS, whereas 1508 proteins with at least one peptide were uniquely quantified by PARADIAS using a library constructed from recovered DIA spectra. We considered that, despite the differences in false discovery rate estimation procedures of benchmarked software, in essence, they all use the similar target-decoy FDR estimation procedure (Reiter *et al*, 2011). This led to the hypothesis that the observed differences between PARADIAS and other methods were due to the way FDR estimation is performed in targeted quantifications. Indeed, by randomly shuffling up to 50% of amino acids in peptide sequences and building *in silico* libraries for targeted approaches (such that none of the shuffled libraries shared precursor fragments with the experimental library, Methods M5), and using these shuffled sequences as search databases to identify the recovered spectra from PARADIAS, up to 20-fold more false identities were reported by other tools when not using a PARADIAS library (Figure 2c, Figure S1-2). A potential explanation is that decoy generation techniques, such as random shuffling, sequence reversal, or introducing specific systematic mutations (Wang *et al*, 2009; Levitsky *et al*, 2017) would generate unrealistic scores, by having very different fragmentation patterns than those present in natural proteomes. The resulting decoy score distributions calculated from DIA data thus allow even 50% mutated peptides to be identified as hits, as opposed to running search engines on deconvolved spectra output from PARADIAS, which uses spectral properties instead of DIA data target-decoy features and results in a more sensitive matching. Therefore, PARADIAS can be used for building high-confidence spectral libraries directly from data, and these can also be used with other targeted approaches to prevent false identification.

Furthemore, we found that PARADIAS-recovered spectra contain on average twice more confidently identified (1% FDR) posttranslationally modified peptides than by using pseudo-spectra from established methods (Table S1-1). Although we consistently identified a total of only 101 modified peptides in our yeast lysate replicates dataset, these were identified directly from a regular chromatography setup, without applying specialized PTM enrichment techniques (Zhao & Jensen, 2009). We demonstrated that recovered spectra can also be *de novo* sequenced, using combinatorial and deep learning approaches (Ma, 2015; Tran *et al*, 2017), resulting in up to 24 times more peptide sequences.

Moving forward, we anticipate that further improvements to the framework will increase not only the quality and quantity of results, but also running time. For this study, the method performed well due to the high-quality, robust chromatographic gradients in our datasets (Vowinckel *et al*, 2018; Navarro *et al*, 2016), i.e. the yeast and HYE110 datasets had less than 5% average variability in retention times. To account for less reproducible gradients, adding a retention time alignment step (Lange *et al*, 2008; Röst *et al*, 2016) would certainly improve spectra recovery and, correspondingly, the number of peptide identifications, as this is crucial for PARAFAC to perform well, since the trilinearity assumption is no longer guaranteed to hold when shifts on the retention time axis are present (Smilde *et al*, 2005). Additionally, a different decomposition method can potentially improve results, i.e. the theoretically better model in this case would be PARAFAC2, which allows for slight nonlinearities in one mode (retention time shifts in this case) (Bro *et al*, 1999), thus alleviating the requirement for robust gradients. However, at the time of this study no efficient implementation existed. Presently, the relative quantification is performed using a library constructed from deconvolved spectra, whereas the sample mode of each component already gives the relative contribution of that component to each sample. In practice, we have seen this to be too imprecise to use for high-quality quantification. Thus, improvements to the decomposition would enable analyte quantification from sample modes directly. Our framework can be readily adapted to other types of DIA data, including sliding MS1 window techniques (Messner *et al*, 2019; Moseley *et al*, 2018) and small molecule metabolomics data (Zhu *et al*, 2014). As the pipeline relies on the high-level Python multiway framework TensorLy (compatible with major machine learning backends (Kossaifi *et al*, 2019)), it can be readily adapted to include additional separation dimensions such ion-mobility separation (d’Atri *et al*, 2018) using either four-way PARAFAC or Tucker3 decomposition. To conclude, as a state-of-the-art computational solution to deconvolve MS scans, PARADIAS shows potential to greatly expand the coverage and efficacy of the DIA approach, and we hope it can serve as a general platform for multiway analysis of mass spectrometry data.

## Methods

### M1. Preprocessing

DIA scans were partitioned and combined to form independent tensors for the decomposition stage. To determine the size of these partitions or “slices”, we used the MS 1 precursor isolation windows or “swaths” to cut the scans along the m/z axis, and a reasonable time window to cut the retention time axis, depending on the chromatography. Partitioning according to swaths is a natural approach, as precursor-product spectra within one swath are independent of those in other swaths. For the yeast replicates and TOR study datasets, the time window was chosen as 1 minute, whereas for the HYE110 dataset, 5 minute windows were taken. This choice balanced the number of expected elutants, due to differences in gradients (e.g. 20 minute vs 40 minutes) and hence, the range of possible models, against the resulting number of slices. To facilitate processing, the scan files were converted to tabular format and partitioned in parallel using the distributed computing framework Apache Spark. Each such slice thus contained the same (m/z, RT) partition for all input scan files, which were then “stacked”, resulting in a (m/z, RT, sample) tensor structure, encoded as a NumPy (Oliphant, 2006) array. Technically, each MS 1 survey scan and its respective MS 2 spectra were aligned along the time axis, as they ought to form a single variable, i.e. the same column in the resulting matrix. The preprocessing step resulted in a collection of independent tensors that span the entire m/z and RT range of each sample. For more details, see Supplementary Note S1-3.

### M2. PARAFAC decomposition

Each slice tensor (**<u>D</u>**) resulting from the preprocessing step was decomposed using PARAFAC into a sample mode *S*, a (retention) time mode *T*, and an m/z mode *M*, plus a residual error term **<u>E</u>**, for a given number of components F spanning a predetermined range. For an explicit form using the Kruskal operator (Kolda & Bader, 2009), see Eq. 1.

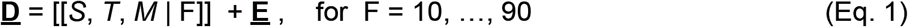

Each of these three mode matrices consist of F components, which correspond to separable analyte signals. That is, assuming perfect decomposition, each column in *S*, each column in *M*, and each row in *T* corresponds to the m/z spectrum, elution profile, and sample contribution of a single peptide. As this number F is unknown *a priori*, we performed the decomposition for an expected number of peptides within a slice. The choice of F value range was informed by inspecting a scan with Spectronaut. A nonnegativity constraint was imposed on all modes. All slice tensors were decomposed in parallel using the TensorLy GPU-adapted implementation. (Kossaifi *et al*, 2019), with PyTorch as a backend.

### M3. Model selection

To select the best model per slice from the range generated in the previous step, we counted the peaks of the time mode of each component of each model, using a continuous wavelet transformation approach (Du *et al*, 2006), implemented in the SciPy package (Virtanen *et al*, 2020). As each analyte should have a single elution peak, we counted, per model, the fraction of components with a single peak. Among all models generated for a slice, we chose the simplest model (lowest F) with maximum fraction of unimodal time modes (see Figure 3c for an illustration). To test the performance of this criterion, we constructed spectra files from two other model sets, with the number of components set to half and twice the optimum F according to unimodality, respectively, for each slice. These spectra files were then analyzed with Crux (Comet and Percolator) and the number of high-confidence peptide identifications was compared. The models with highest unimodality had the best performance (Figure 3b).

### M4. Identification and Quantification

The spectra from the best models are saved to an mzXML file. This resembles a DDA file since each scan entry consists of the MS 2 part of the deconvolved spectrum, along with its corresponding highest intensity MS 1 peak as precursor. This file was then searched using Comet and MS-GF+ to produce PSMs in conjunction with a proteome FASTA database, using the same mass tolerance as the initial acquisition (i.e. 40 ppm for the yeast replicates and TOR study datasets, and 50 ppm for the HYE110 dataset). Comet results were then filtered using Percolator at 1% FDR. The confidence assessment for both MS-GF+ and Percolator was done using reversed decoys. To search for PTMs, MS-GF+ was pre-configured to account for acetylations, succinylations, phosphorylations, and core 1 GalNAc glycosylations as variable modifications anywhere in the peptide, allowing for maximum 384 variable modifications.

The output PARADIAS mzXML file was used to construct a spectral library following the protocol in (Schubert *et al*, 2015), using Comet as a source of PSMs. DIA-NN was then run on the initial DIA scan files using this library to produce peptide quantities, using default parameters. Benchmark results for the HYE110 dataset were obtained using the lfqbench R package (Navarro *et al*, 2016).

### M5. False positive assessment

Proteome FASTA database files were randomized such that 50% and 30% of each trypsin-digested protein sequence got shuffled, for the yeast replicates and HYE110 results, respectively. DIA-NN and Spectronaut were run in library-free mode (called “directDIA” for the latter), generating a spectral library *in silico* based on the shuffled database. The same mode was used for their baseline results on the original (nonrandomized) databases. As Skyline does not have *in silico* library generation, the library created by DIA-NN from shuffled sequences was used as its source of transitions, and published spectral libraries (Vowinckel *et al*, 2018; Navarro *et al*, 2016) for baseline results. To assess the effect of using PARADIAS as a preprocessor, all tools were run using a spectral library created from the PARADIAS deconvolved spectra and the corresponding shuffled library, following the protocol in (Schubert *et al*, 2015). For all these runs, we removed any resulting spectra that were also found in the published libraries, thus ensuring we generated exclusively false positives. In terms of parameters, Skyline was run analogously with (Navarro *et al*, 2016), Spectronaut and DIA-NN with default settings, except with a 100% FDR threshold to allow selecting results at different false discovery levels.

### M6. Scan reconstruction from a PARAFAC model

The output PARAFAC sample mode *S*, retention time (RT) mode *T*, and m/z mode *M*, consisting of F components resulting from the decomposition of a dataset, were used to reconstruct a pseudo-scan comprised of the deconvolved analytes. A pseudo-scan *P*_*i*_ is an (m/z, RT) matrix corresponding to the input DIA scan *i* in the dataset. It is obtained by summing over the outer products of the m/z mode *m* and RT mode *t*, multiplied by contribution to scan *i* from the sample mode *s*, for all unimodal PARAFAC components *r* in the model.

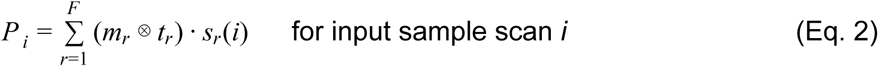

Lastly, the resulting intensities in *P*_*i*_ were scaled back to the values in the corresponding scan *i*, and multiplied by the model R^2^, as PARAFAC solutions do not preserve the scaling of the input tensor (Smilde *et al*, 2005). This is, in effect, the reverse operation to PARAFAC for a single input sample, discarding the residuals (Eq. 1 and Note S1-1, Eq. 2). The above operation was done piecewise, for each independent tensor produced by PARADIAS partitioning of the input dataset.

### M7. *De novo* sequencing

The output PARADIAS mzXML was converted to MGF format and subsequently set as input to Novor (Ma, 2015) and DeepNovo (Tran *et al*, 2017). Novor was run using a mass tolerance of 50 ppm and DeepNovo a tolerance of 10 ppm. Novor was set to CID fragmentation and TOF mass analyzer, using otherwise default parameters. For DeepNovo, the pre-trained *yeast.low.coon_2013* model was used, with a beam size of 5. For the baseline DIA-Umpire results, only the highest quality (Q1) extracted features were used, since these are far more likely to lead to good sequencing results, as good fragment coverage is needed (Muth & Renard, 2018). Moreover, we considered only sequences that appear in at least 6 out of 9 replicates based on DIA-Umpire features.

### M6. Software availability

The PARADIAS pipeline is available on GitHub at https://github.com/fburic/paradias.

## Acknowledgements

The computations were mainly performed on resources at Chalmers Centre for Computational Science and Engineering (C3SE), and partially at High Performance Computing Center North (HPC2N), provided by the Swedish National Infrastructure for Computing (SNIC). Mikael Öhman at C3SE is acknowledged for assistance concerning technical aspects in making the code run on the C3SE resources. We thank Hieu Tran for assistance in running DeepNovo and Vadim Demichev for assistance in running DIA-NN. We also thank Kate Campbell for useful discussions and early feedback on the manuscript. JZ, FB, and AZ were supported by SciLifeLab funding.

