## Supplementary Material for "Parallel factor analysis enables quantification and identification of highly-convolved data independent-acquired protein spectra"

#### Table of Contents

Supplementary Figures - p. 2 - 6
Supplementary Tables - p. 7 - 12
Supplementary Notes - p. 13 - 24
Supplementary References - p. 25

### Supplementary Figures

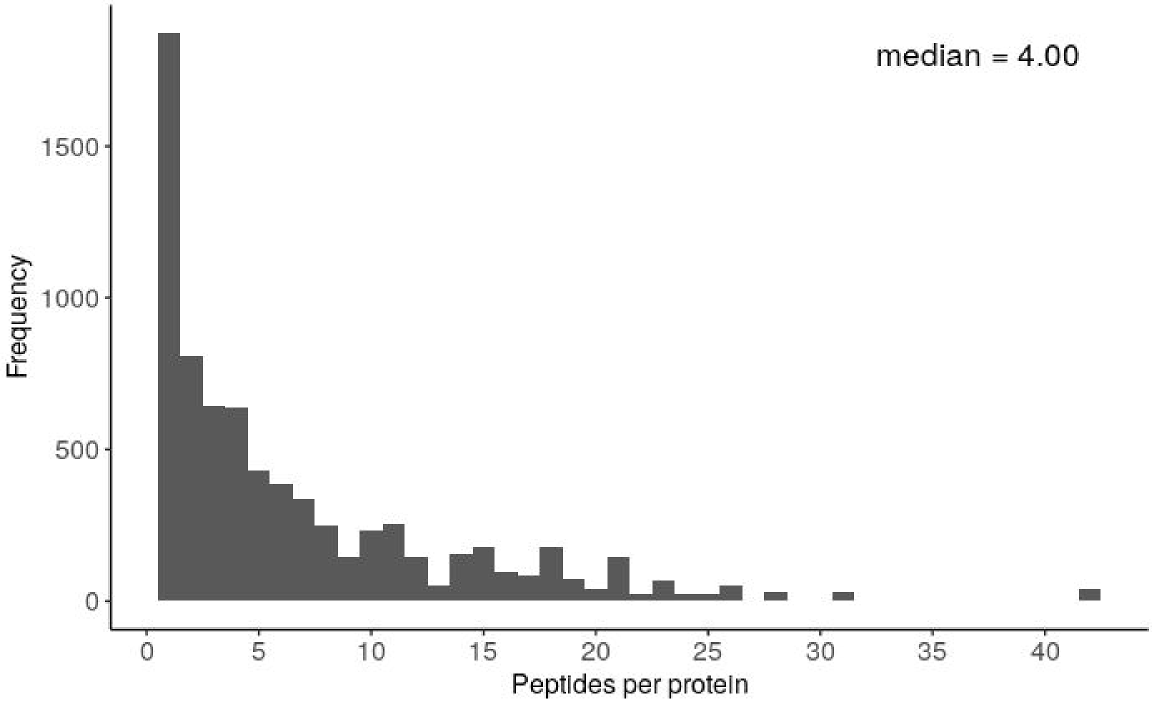

**Figure S1-1.** Number of identified peptides per protein for the HYE110 dataset. A total of 5908 peptides were identified by Crux and MS-GF+ at FDR 1% on PARADIAS output.

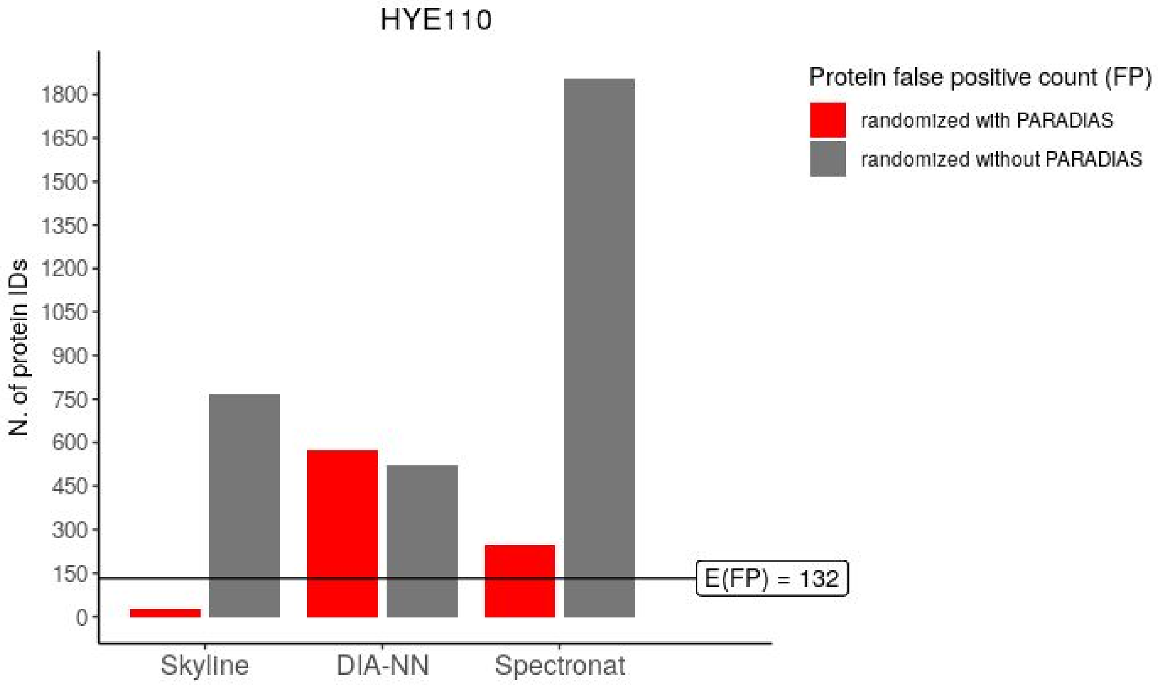

**Figure S1-2. False positives count underestimation on the HYE110 dataset, across different software, at 5% FDR.** The bar plots show the number of protein IDs under a randomized trial (where all corresponding peptides are false positives), both with and without using PARADIAS as a source of the spectral library, compared to the average expected number of false positives based on baseline runs for each tool. The results are presented as false protein identifications to demonstrate the impact on the total proteome, i.e. the protein considered as false was identified if at least one of its randomized peptides was quantified in the sample.

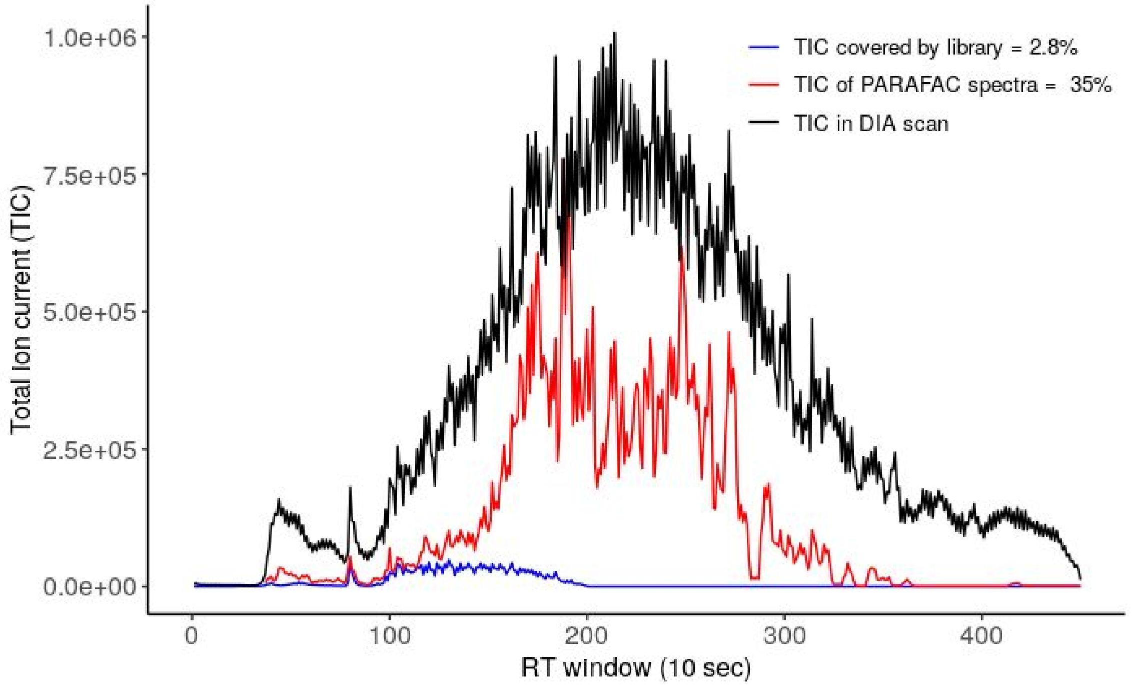

**Figure S1-3.** Typical DDA spectral library coverage (2.8%) of the total ion current (TIC) in a centroided DIA scan, compared with that of recovered PARAFAC components (35%). Shown is the TIC per 1 second window of one yeast lysate replicate (gray), along with the TIC of the scan spectra matched to the library (blue), and the TIC of a pseudo-scan reconstructed from PARAFAC output (red). Allowing for a 5-minute retention time (RT) and a 40 ppm m/z tolerance, the spectral library matched 29.53% of the m/z space in the DIA scan. At least 4 library fragments (product m/z) needed to match for a spectrum to be considered covered. The reconstructed PARAFAC pseudo-scan was obtained by taking the sum of the outer products of each component’s m/z and time mode, scaled according to the sample mode coefficient for this scan and the maximum scan intensity.

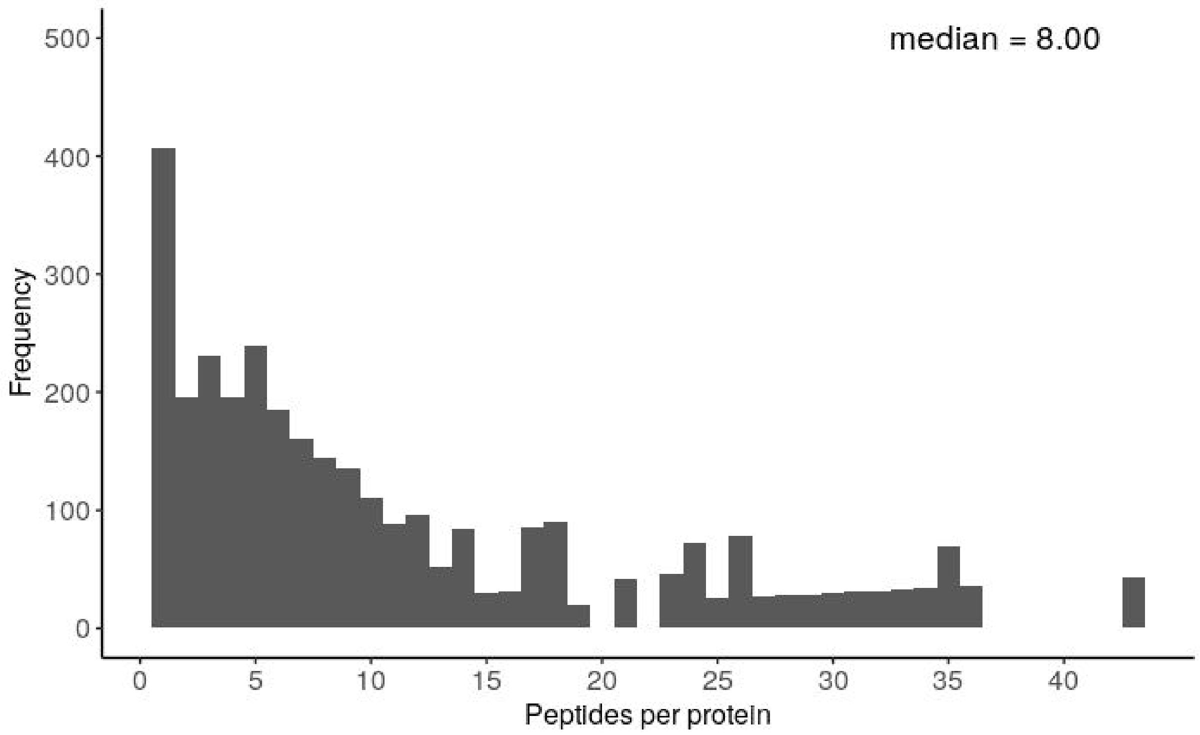

**Figure S1-4.** Number of identified peptides per protein for the yeast replicates dataset. A total of 2653 peptides (840 proteins) were identified by Crux and MS-GF+ at FDR 1% on PARADIAS output.

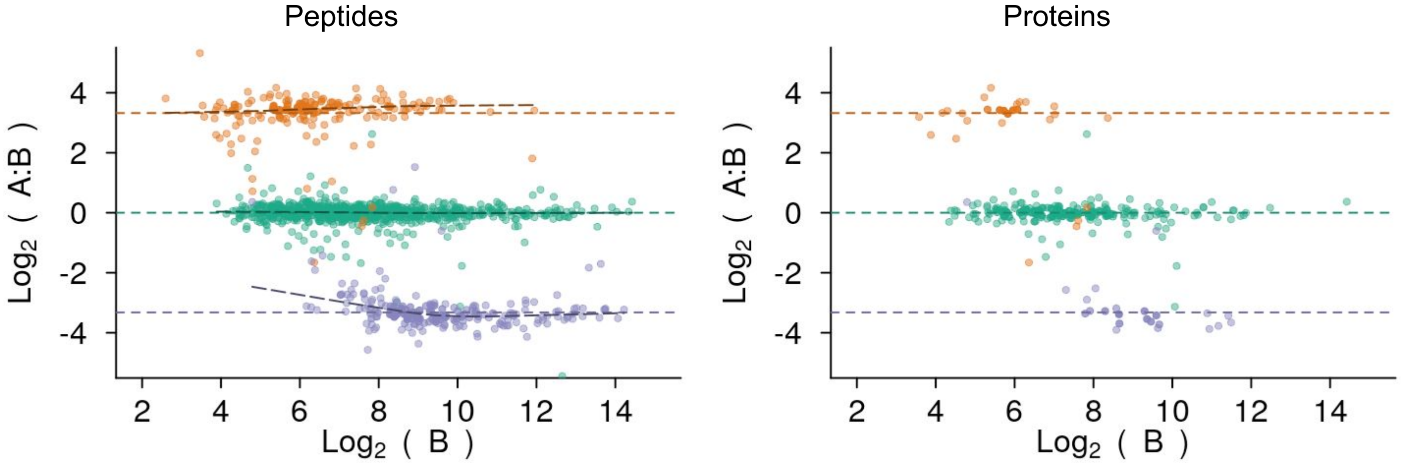

**Figure S1-5.** Peptide- and protein-level LFQbench results on the HYE110 dataset, using a library constructed from PARADIAS output spectra and quantification performed by DIA-NN. The dataset contains two sample mixtures A and B, with known composition, namely A with 67% human (green), 30% *S. cerevisiae* (orange), and 3% *E.coli* (purple), and B with 67% human, 3% *S. cerevisiae*, and 30% *E.coli*). The plots show the spread of quantity ratios per organism, as well as regression lines per organism (dashed black lines), against the expected ratios (horizontal dashed lines with colors matching the organism). After filtering DIA-NN results with an FDR threshold of 1% and subsequently applying the standard LFQbench criteria for valid results, 3590 peptides and 244 protein ratios remained. Lower-intensity values exhibit lower quality (higher variability).

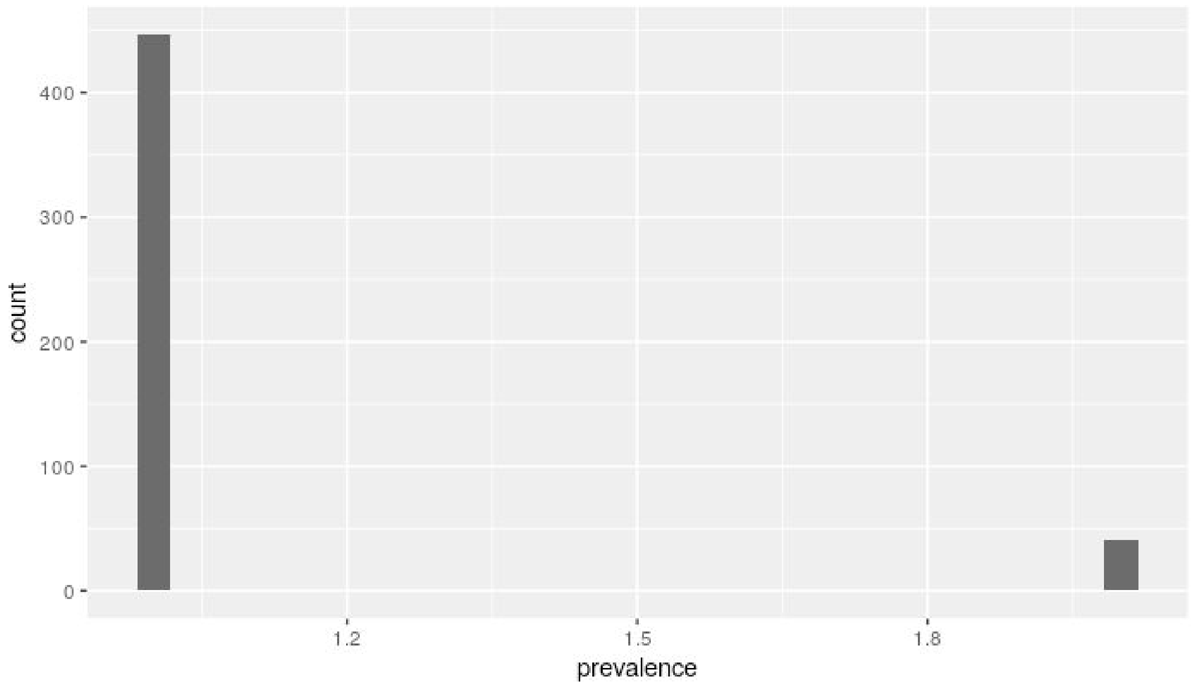

**Figure S1-6.** Prevalence (presence in technical replicates) of n = 487 peptide sequences found as modified by MS-GF+ in DIA-Umpire output pseudo-spectra files. The sequences were stripped of the modifications to allow for variation between replicates. 40 of these peptides appear in 2 replicates, while all the rest only appear in 1 replicate.

### Supplementary Tables

| | PARADIAS / MS-GF+ | DIA-Umpire / MS-GF+ (median across replicates) |
|---|---|---|
| Acetylations | 108 | 47 |
| Phosphorylations | 26 | 13 |
| Succinylation | 30 | 21 |
| GalNAc (core 1) glycosylations | 22 | 14 |
| Modified peptides | 101 | 56 |
| ***Run time*** | *11.5 h* | *70 h* |

**Table S1-1. Post-translational modification counts found with MS-GF+ on PARADIAS and DIA-Umpire outputs at 1% FDR.** For the latter, median PTM counts across the replicates is given. All three DIA-Umpire output files, of decreasing quality (Q1, Q2, and Q3), were considered for each technical replicate.

| protein_id | peptide_sequence |
|---|---|
| sp\|P10127\|ADH4_YEAST | K.DY(UniMod:21)K(UniMod:64)K(UniMod:64)ALIVTDPGIAAIGLSGR.V |
| sp\|P0C0W9\|RL11A_YEAST | R.LTRASK(UniMod:64)VLEQLSGQTPVQSK.A |
| sp\|P39954\|SAHH_YEAST | R.LTK(UniMod:64)LS(UniMod:21)KVQSEYLGIPEEGPFK.A |
| sp\|P39954\|SAHH_YEAST | R.LT(UniMod:21)K(UniMod:64)LSKVQSEYLGIPEEGPFK.A |
| sp\|P00330\|ADH1_YEAST | K.S(UniMod:X)PIK(UniMod:64)VVGLSTLPEIYEK.M |
| sp\|P02829\|HSP82_YEAST | R.Y(UniMod:1)NS(UniMod:1)T(UniMod:21)K(UniMod:1)SVDELTSLTDYVTR.M |
| sp\|P53723\|YN8B_YEAST | M.SSS(UniMod:1)IFGPLTGFLER.V |
| sp\|P04801\|SYTC_YEAST | M.S(UniMod:1)ASEAGVTEQVKK.L |
| sp\|P04801\|SYTC_YEAST | M.SA(UniMod:1)SEAGVTEQVKK.L |
| sp\|P22217\|TRX1_YEAST | K.EVA(UniMod:1)KVVGANPAAIK.Q |
| sp\|P53723\|YN8B_YEAST | M.SS(UniMod:1)SIFGPLTGFLER.V |
| sp\|P53723\|YN8B_YEAST | M.S(UniMod:1)SSIFGPLTGFLER.V |
| sp\|P22217\|TRX1_YEAST | K.EVAK(UniMod:1)VVGANPAAIK.Q |
| sp\|P32590\|HSP79_YEAST | M.S(UniMod:1)TPFGLDLGNNNSVLAVAR.N |
| sp\|P14020\|DPM1_YEAST | M.S(UniMod:1)IEYSVIVPAYHEK.L |

| | |
|---|---|
| sp\|P00549\|KPYK1_YEAST | R.AEVSDVGNAILDGADCVMLS(UniMod:21)GETAK.G |
| sp\|P11484\|SSB1_YEAST | K.DA(UniMod:1)KIS(UniMod:1)KSQIDEVVLVGGSTR.I |
| sp\|P36105\|RL14A_YEAST | K.LA(UniMod:1)AIVEIIDQKK.V |
| sp\|Q12499\|NOP58_YEAST | M.AY(UniMod:1)VLTETSAGYALLK.A |
| sp\|Q12499\|NOP58_YEAST | M.A(UniMod:1)YVLTETSAGYALLK.A |
| sp\|P07149\|FAS1_YEAST | R.LNA(UniMod:1)DFQKPWFATVNGQAR.D |
| sp\|P25443\|RS2_YEAST | M.S(UniMod:1)APEAQQQK.R |
| sp\|P0CX48\|RS11B_YEAST | M.S(UniMod:1)TELTVQSER.A |
| sp\|P38113\|ADH5_YEAST | R.A(UniMod:1)DTREALDFFAR.G |
| sp\|P04147\|PABP_YEAST | M.A(UniMod:1)DITDKTAEQLENLNIQDDQK.Q |
| sp\|P46672\|ARC1_YEAST | M.S(UniMod:1)DLVTKFESLIISKYPVSFTK.E |
| sp\|P16862\|PFKA2_YEAST | K.VHS(UniMod:21)YTDLAYR.M |
| sp\|P32565\|RPN2_YEAST | M.S(UniMod:1)LTTAAPLLALLR.E |
| sp\|P38625\|GUAA_YEAST | K.VA(UniMod:1)S(UniMod:21)RIVNEVDGVAR.V |
| sp\|P26637\|SYLC_YEAST | M.S(UniMod:1)SGLVLENTAR.R |
| sp\|P16550\|APA1_YEAST | M.S(UniMod:1)IPADIASLISDKYK.S |
| sp\|P39954\|SAHH_YEAST | M.S(UniMod:1)APAQNYK.I |
| sp\|P00331\|ADH2_YEAST | K.EELFTS(UniMod:1)LGGEVFIDFTK.E |
| sp\|P40075\|SCS2_YEAST | M.S(UniMod:1)AVEISPDVLVYK.S |
| sp\|P00549\|KPYK1_YEAST | R.AEVS(UniMod:21)DVGNAILDGADCVMLSGETAK.G |
| sp\|P38011\|GBLP_YEAST | M.A(UniMod:1)SNEVLVLR.G |
| sp\|P06169\|PDC1_YEAST | M.S(UniMod:1)EITLGKYLFER.L |
| sp\|Q12165\|ATPD_YEAST | K.RSY(UniMod:21)AEAAAASSGLK.L |
| sp\|P36008\|EF1G2_YEAST | M.S(UniMod:1)QGTLYINR.S |
| sp\|P00958\|SYMC_YEAST | K.CGA(UniMod:1)LLDPFELINPR.C |
| sp\|P53277\|SYF2_YEAST | K.KRY(UniMod:1)EARK(UniMod:64)R.Q |
| sp\|Q06142\|IMB1_YEAST | M.S(UniMod:1)TAEFAQLLENSILSPDQNIR.L |
| sp\|P16550\|APA1_YEAST | M.S(UniMod:1)IPADIASLISDK.Y |
| sp\|P02994\|EF1A_YEAST | R.RSGK(UniMod:1)KLEDHPK.F |
| sp\|P36534\|RM40_YEAST | R.VVVMS(UniMod:1)GA(UniMod:1)S(UniMod:1)K(UniMod:64).G |
| sp\|P40075\|SCS2_YEAST | M.SA(UniMod:1)VEISPDVLVYK.S |
| sp\|P06367\|RS14A_YEAST | M.S(UniMod:1)NVVQAR.D |
| sp\|P38867\|YHX7_YEAST | R.K(UniMod:1)YFY(UniMod:1)TGLLK(UniMod:64)K.T |
| sp\|P00958\|SYMC_YEAST | M.S(UniMod:1)FLISFDK.S |
| sp\|Q12074\|SPEE_YEAST | M.A(UniMod:1)QEITHPTIVDGWFR.E |
| sp\|P42947\|YJK7_YEAST | K.SIS(UniMod:X)DLGISNNDNK.N |
| sp\|P40319\|ELO3_YEAST | K.T(UniMod:X)VK(UniMod:1)K(UniMod:1)ES(UniMod:1)EVSGSVASGSSTGVK.T |
| sp\|P25627\|BUD23_YEAST | K.VAK(UniMod:1)DS(UniMod:1)K(UniMod:64)FTGR.K |

| | |
|---|---|
| sp\|P40151\|WRIP1_YEAST | K.QVWNIENPLKLS(UniMod:1)R.S |
| sp\|P23542\|AATC_YEAST | M.S(UniMod:1)ATLFNNIELLPPDALFGIK.Q |
| sp\|P23542\|AATC_YEAST | M.SA(UniMod:1)TLFNNIELLPPDALFGIK.Q |
| sp\|Q01477\|UBP3_YEAST | K.S(UniMod:X)WS(UniMod:1)A(UniMod:1)IASDAIK.S |
| sp\|P38088\|SYG_YEAST | K.IK(UniMod:64)RMSVEDIK(UniMod:64)K.A |
| sp\|Q04839\|GFD1_YEAST | K.QK(UniMod:1)PS(UniMod:1)HK(UniMod:64)R.S |
| sp\|P07246\|ADH3_YEAST | R.Y(UniMod:21)VVDTSK.- |
| sp\|P20081\|FKBP_YEAST | M.S(UniMod:1)EVIEGNVK.I |
| sp\|P38130\|KTR3_YEAST | K.Y(UniMod:1)FSAGNY(UniMod:1)K.L |
| sp\|P40046\|VTC1_YEAST | M.S(UniMod:1)SAPLLQR.T |
| sp\|P49626\|RL4B_YEAST | K.K(UniMod:64)AEKT(UniMod:21)GT(UniMod:21)KPAAVFAETLK.H |
| sp\|P49626\|RL4B_YEAST | K.KAEK(UniMod:64)T(UniMod:21)GT(UniMod:21)KPAAVFAETLK.H |
| sp\|Q06409\|YL422_YEAST | K.FYNGLS(UniMod:1)VA(UniMod:1)NK.A |
| sp\|Q06116\|YP117_YEAST | K.K(UniMod:1)T(UniMod:X)FNWS(UniMod:21)LK(UniMod:1)LRMK.D |
| sp\|P49089\|ASNS1_YEAST | R.IPST(UniMod:X)PIDYMAIR.H |
| sp\|Q03652\|DML1_YEAST | K.IFLNSVVDK(UniMod:1)VS(UniMod:21)K.T |
| sp\|P53101\|STR3_YEAST | R.ENGK(UniMod:64)YLFNK(UniMod:1)LNK.N |
| sp\|P53246\|ENV11_YEAST | R.SFK(UniMod:64)FT(UniMod:X)K(UniMod:64)SK.K |
| sp\|P23301\|IF5A1_YEAST | K.S(UniMod:1)RPCK(UniMod:1)IVDMSTSK.T |
| sp\|P00958\|SYMC_YEAST | M.S(UniMod:X)FLISFDKSKK.H |
| sp\|Q03735\|NAB6_YEAST | R.FY(UniMod:21)K(UniMod:64)K(UniMod:1)VK(UniMod:1)R.P |
| sp\|P24004\|PEX1_YEAST | -.MT(UniMod:X)TTK(UniMod:64)RLK.F |
| sp\|P09201\|F16P_YEAST | K.K(UniMod:64)S(UniMod:1)PNGK(UniMod:64)LR.L |
| sp\|P38266\|AIM3_YEAST | R.NFS(UniMod:1)LK(UniMod:1)ANEY(UniMod:21)PK.E |
| sp\|Q07915\|RLP24_YEAST | K.WTK(UniMod:1)A(UniMod:1)FR.K |
| sp\|P25567\|SRO9_YEAST | M.SA(UniMod:1)ETAAANTATAPVPEVQEQESSK.S |
| sp\|P25567\|SRO9_YEAST | M.S(UniMod:1)AETAAANTATAPVPEVQEQESSK.S |
| sp\|P50276\|MUP1_YEAST | K.NGGEKNY(UniMod:1)LEA(UniMod:1)IFRK.P |
| sp\|Q03780\|YD239_YEAST | R.K(UniMod:1)PS(UniMod:1)VPS(UniMod:1)TIKK.S |
| sp\|P32465\|HXT1_YEAST | K.S(UniMod:X)A(UniMod:1)S(UniMod:21)WVPVS(UniMod:21)KR.G |
| sp\|P32465\|HXT1_YEAST | K.S(UniMod:21)A(UniMod:1)S(UniMod:X)WVPVS(UniMod:21)KR.G |
| sp\|P16521\|EF3A_YEAST | K.K(UniMod:1)A(UniMod:1)K(UniMod:64)DILDEFRK.R |
| sp\|P15019\|TAL1_YEAST | M.S(UniMod:1)EPAQKK.Q |
| sp\|Q08226\|CRT10_YEAST | K.GY(UniMod:1)AT(UniMod:21)LY(UniMod:1)VASR.G |
| sp\|P11972\|SST2_YEAST | K.YT(UniMod:X)FT(UniMod:X)T(UniMod:X)K.A |
| sp\|P32190\|GLPK_YEAST | K.YWEVAVERS(UniMod:1)K.G |
| sp\|P28742\|KIP1_YEAST | R.T(UniMod:X)AQFEA(UniMod:1)NK.R |
| sp\|P16474\|BIP_YEAST | K.KK(UniMod:64)HGIDVS(UniMod:1)DNNK(UniMod:64).A |

| | |
|---|---|
| sp\|P10962\|MAK16_YEAST | K.T(UniMod:X)PERA(UniMod:1)HTPA(UniMod:1)K.L |
| sp\|P53336\|YG5X_YEAST | K.DK(UniMod:1)K(UniMod:1)HEK(UniMod:64)K(UniMod:64)SR.S |
| sp\|Q05670\|FUS2_YEAST | K.S(UniMod:X)RY(UniMod:1)MT(UniMod:X)K(UniMod:1).R |
| sp\|P42945\|UTP10_YEAST | R.K(UniMod:1)LT(UniMod:21)IILEALDK.V |
| sp\|P47024\|YJ41B_YEAST | K.TQK(UniMod:64)RS(UniMod:X)NK(UniMod:64)VYNSK.K |
| sp\|Q03516\|RSN1_YEAST | R.FLK(UniMod:64)PHIYY(UniMod:1)S(UniMod:21)YK.A |
| sp\|Q04089\|DOT1_YEAST | K.K(UniMod:1)S(UniMod:X)S(UniMod:21)T(UniMod:X)TTK.K |
| sp\|P00331\|ADH2_YEAST | M.S(UniMod:1)IPETQK.A |
| sp\|P00330\|ADH1_YEAST | M.S(UniMod:1)IPETQK.G |
| sp\|P32660\|ATC5_YEAST | R.FT(UniMod:X)Y(UniMod:1)DS(UniMod:1)FQK.F |

**Table S1-2. List of PTMs found in the yeast replicate dataset.** Unimod accessions numbers: 1 = Acetylation, 21 = Phosphorylation, 64 = Phosphorylation, X = stand-in for GalNAc glycosylation for uniform notation (This PTM is not present in UniMod)

| Dataset | Sample | Description |
|---|---|---|
| 1 | Yeast lysate (Vowinckel et al. 2018) | 9 technical replicates |
| 2 | Yeast lysate (Vowinckel et al. 2018) | 30 samples (excluding replicates), TOR study |
| 3 | Human, yeast, and *E.coli* lysate (Navarro et al. 2016) | LFQ bench HYE110, with 64 variable width precursor isolation windows |

**Table S1-3.** Datasets Included in this study.

| Parameter (YAML variable) | Standard Value | Description |
|---|---|---|
| `window_size_sec` | 60 | How wide the RT windows should be (in seconds), when slicing the data set into (m/z, RT) slices. |
| `avg_peak_fwhm_sec` | 12 | Expected FWHM (full width at half maximum) of an analyte's elution profile (in seconds), assuming it has an approximate Gaussian shape. Used only to inform the peak detection procedure and needs not be highly accurate. |
| `library` | Relative path of FASTA file | Protein FASTA database file name. The term "library" is due to some tools like Comet referring to this file as such. |
| `decoy_library` | Relative path of FASTA file | Protein decoys FASTA DB file name. |
| `isobaric_mixed_library` | Relative path of FASTA file | Mixed protein target and decoy database in which all instances of the amino acid I have been replaced with L, since these have equal mass. Used for exact tag matching of *de novo* sequences. |
| `decoy_prefix` | "decoy_" | Protein ID prefix to mark it as a decoy in its FASTA database. |
| `parafac_init` | "random" | PARAFAC initialization ("first guess") method, as implemented in TensorLy. "svd" may also be used, usually with better results, but this risks exhausting memory, given it's quadratic space complexity as a function of the longest dimension of the tensor. |
| `parafac_max_iter` | 5000 | Maximum number of PARAFAC fitting iterations. |
| `parafac_min_comp` | 10 | Start of the inclusive range of number of PARAFAC components to decompose for. |
| `parafac_max_comp` | 90 | End of the range of number of components. |
| `parafac_backend` | "pytorch" | The tensor framework to be used by TensorLy for the PARAFAC algorithm. Currently only "pytorch" is fully integrated. |
| `parafac_avail_ram_gb` | 16 | Amount of GPU RAM (in GB). |
| `analysis_pipeline` | "crux" | The identification tool to use. Currently only "crux" and "msgf+" are supported. |
| `comet_mass_tol_ppm` | 40 | The mass tolerance of the SWATH scans. |
| `msgf_modifications` | "path/to/msgf_mods.txt" | File name of the text file specifying which post-translational modifications MS-GF+ should search for. See MS-GF+ documentation for details. |

| | | |
|---|---|---|
| `msgf_threads` | 20 | MS-GF+ searches are parallelized. Value should be set to the number of CPUs |
| `model_index` | "path/model_index.feather" | Tabular file in Feather format that indexes all models generated by PARADIAS across all slices. Each model has a unique ID across the dataset. |
| `spectrum_index` | "path/spectrum_index.feather" | Tabular file in Feather format that indexes all spectra (mass modes), for all models (and components) generated by PARADIAS across all slices. Each spectrum has a unique ID across the dataset. |
| `time_modes_values` | "path/time_mode_values_all_models.feather" | Tabular file in Feather format that collects peak counts for all time modes (in all components), for all models generated by PARADIAS across all slices. |
| `intensity_lower_percentage_cutoff` | 1 | Percentile to use for baseline noise filtering of output PARADIAS spectra. All values within this bin are removed. |
| `percolator_fdr` | 0.01 | FDR threshold to be used by Percolator to filter PSMs from Comet. Normally set slightly more permissive at this stage, to include more candidate results, for downstream analyses to work with. |
| `quant_lib_mayu_fdr` | 0.1 | Set higher to be permissive. The quantification software does its own FDR estimation and filtering at 1% |
| `lower_mz_frag` | 100 | Start of the fragment m/z range to be used in the library construction. |
| `upper_mz_frag` | 2000 | End of the fragment m/z range to be used in the library construction. |
| `quant_library_spectrast_max_frag_annot_err` | 0.05 | Maximum error allowed at the annotation of a fragment ion. |
| `sequencer` | "novor" | Which *de novo* sequencing tool to use. Currently integrated: "novor". |

**Table S1-4**. Description and standard values for important pipeline parameters.

### Supplementary Notes

#### Note S1-1. The PARAFAC procedure

Parallel factor analysis (PARAFAC) (Harshman 1970), also known as canonical decomposition (CANDECOMP) (Carroll, Douglas Carroll, and Chang 1970), is a procedure for decomposing tensors (multidimensional arrays) into products of linearly contributing factors or “modes” (Figure S1-1-1a, Eq. 1). It can be seen as a higher-dimensional generalization of singular value decomposition (SVD) and principal component analysis (PCA). Unlike these, however, a PARAFAC decomposition is unique (except for trivial permutation and scaling of the component matrices) for a given number F of components in each factor (Smilde, Bro, and Geladi 2005). Moreover, the resulting mode values have a natural interpretation, as they directly represent the contribution of each e.g. analyte in the original data (Smilde, Bro, and Geladi 2005). There is an abundance of literature on such methods. We refer the reader to a general mathematical introduction in (Kolda and Bader 2009) and a more applied treatment in (Smilde, Bro, and Geladi 2005), which delves into other aspects such as data preprocessing. When applied to LC-MS data, the assumption is that each separable analyte (peptides in our case) corresponds to a PARAFAC component (Figure S1-1-1b, Eq. 2) (Skov and Bro 2008). Analytes that have identical mass spectra and elution curves would not be separable (Johnsen, Amigo, and Skov 2014).

| | |
|---|---|
| **a** 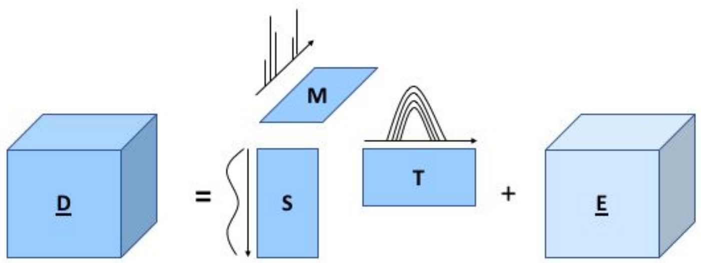  | $\underline{\mathbf{D}} = [\![S, T, M]\!] + \underline{\mathbf{E}}$ (Eq. 1) <br> **D** is the tensor product of the sample mode *S*, the (retention) time mode *T*, and the mass mode *M*, plus the error term **E**. |
| **b** 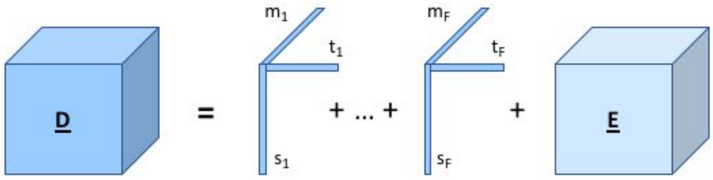  | $\underline{\mathbf{D}} = \sum_{r=1}^{F} s_r \otimes t_r \otimes m_r + \underline{\mathbf{E}}$ (Eq. 2) <br> where $\otimes$ denotes the outer product of the three vectors $s_r$, $t_r$, and $m_r$, associated to the sample, time, and mass contribution of each component r, respectively. |

**Figure S1-1-1**. **Two equivalent ways to express the PARAFAC decomposition of the data tensor D**. The tensor consists of stacked SWATH LC-MS/MS maps, which are decomposed into **a**) a trilinear combination of three matrices or “modes”, corresponding to m/z spectra (the mass mode **M**), elution time (the time mode **T**), and sample variation (the sample mode **S**), plus a residual error term **E**. These matrices span F components, which correspond to separable peptide signals. That is, assuming perfect decomposition, each column in **S**, each column in **M**, and each row in **T** corresponds to the m/z spectrum, elution profile, and sample contribution of a single peptide. The second formulation in **b**) puts this into focus, by expressing the data tensor as the sum of *F* individual peptide contributions along the mass, time, and sample dimensions. This outer product of the three mode vectors associated with a peptide is called a component.

#### Model selection

A critical step in constructing a PARAFAC model is deciding *a priori* the number of components F. Setting F equal to the rank of the tensor, i.e. the smallest number of components that sum to form the tensor, is a natural choice. It is however known that computing the rank of a tensor is NP-complete (Håstad 1989), which implies there are no algorithms that can find the correct rank in a reasonable amount of time. Slightly overestimating F may not be so problematic, since the extra few components may be informative and there may be more than one valid model (Johnsen, Amigo, and Skov 2014). On the other hand, underestimating F leads to improper separation of components. Also problematic is the fact that PARAFAC models do not “nest”, i.e. a model for F+1 is not simply a model for F with an extra component (Smilde, Bro, and Geladi 2005). Deciding the value automatically is, in general, an open problem (Liu et al. 2016), and the various diagnostics and procedures (Bro and Kiers 2003) used to this end often require human verification, which is not feasible for fully automated, high-throughput workflows in which hundreds of thousands of models need to be examined. Our approach is to generate all models within a reasonable F range (estimated by inspecting the scans with tools such as Spectronaut), then select the best models in a subsequent step. We found that simply imposing non-negativity constraints on all modes during decomposition, then selecting best models based only on the unimodality of each component's time mode (as each analyte should have a single elution peak) yields good results with downstream tools.

#### Algorithm 1. Best Model Selection

**Procedure Select_Best_Model**:

```
For each Slice(swath, RT window),
      For each Model(F), with F = F_lower .. F_upper,
            For each component k = 1 .. F,
                  peak_count(k) <- Time_Mode_Peak_Count(Model(F), k)
            n_unimodal_components <- |{k : peak_count(k) == 1 }|
            unimodal_fraction <- n_unimodal_components / F
      Best_Model <- Model(argmax {k = 1 .. F}{unimodal_fraction})
      # first max val considered
```

**Function Time_Mode_Peak_Count**:

```
avg_peak_window_frac <- avg_peak_fwhm_sec / window_size_sec
expected_peak_width_scaled_to_data <- length(time_mode) * avg_peak_window_frac

null_threshold <- 0.1 * max(elution_profile)
elution_profile[elution_profile <= null_threshold] <- 0

peaks <- scipy.signal.find_peaks_cwt(
      elution_profile,
      widths=np.arange(1,expected_peak_width_scaled_to_data * 2)
      )
```

**Parameters**

| | |
|---|---|
| `F_lower, F_upper` | Lower and upper value for the number of components *F* to decompose for (across all slices) |
| `window_size_sec` | RT window size [s] used to slice input data tensor |
| `avg_peak_fwhm_sec` | The expected peak FWHM, to inform the peak finding procedure. This was estimated manually, by opening a scan in Spectronaut and determining the base width of iRT peptides.<br>This width was considered as FTWM of a Gaussian, so therefore the FWHM = FWTM sqrt(ln2 / ln10) |

| | |
|---|---|
| `null_threshold` | Value below which elution profile values are set to zero to help peak detection. (A form of baseline noise filtering.)<br>Set to 0.1 of the max elution profile value throughout this study. |

#### Note S1-2. *De novo* peptide sequencing

The task of deducing the constituent amino acids and their correct ordering in the peptide chain from MS spectra is referred to as *de novo* sequencing. This approach is especially useful for measuring the proteome of organisms that have incomplete protein databases or for improving the quality and speed of library searches (Muth and Renard 2018). The underlying combinatorial problem, however, grows exponentially with the peptide length, combined with the fact that real spectra can be much noisier than their theoretical counterparts, which makes this a very difficult problem (Muth and Renard 2018). Such methods are thus expected to benefit from spectral deconvolution.

We used DeepNovo (Tran et al. 2017), which employs deep learning architectures trained on different organisms to translate spectra into amino acid sequences, and the established Novor software (Ma 2015), which uses decision trees built using machine learning from a NIST human peptide spectral library. We compared the performance of DeepNovo on PARAFAC output with that on features extracted by DIA-Umpire from the SWTH scans, and considered only sequence score (probability of correctness) above 80%, and, for the DIA-Umpire results, sequences that appear in at least 6 out of 9 replicates. For DIA-Umpire, only the highest quality (Q1) extracted features were used, since these are far more likely to lead to good sequencing results, as good fragment coverage is needed (Muth and Renard 2018). DeepNovo yielded 415 peptide sequences from PARADIAS output and 101 from DIA-Umpire, while Novor yielded 407 sequences from PARADIAS output and 17 from DIA-Umpire.

### Note S1-3. Preprocessing steps

The input DIA scan files are processed as below by the PARADIAS pipeline. For more information regarding software versions and parameters, see the Software Versions section.

1. Scan files are converted to mzML using *msconvert*, with peak picking on MS both levels
2. Scan files are converted from mzML to CSV with the following constraints:
    a. intensity values below 1 are discarded
    b. 10 decimals for m/z
    c. 4 decimals for retention time (RT)
    d. 8 decimals for intensity
3. Swath windows are adjusted to avoid overlaps: the adjusted window end is set to the midpoint of the unadjusted window end and the next window's unadjusted window start. E.g. the isolation windows (400, 416) and (415, 432) become (400, 415.5) and (415.5, 431.6).
4. Samples are split into (swath, RT window) slices using Apache Spark
5. Each slice is converted into a tensor (3-dimensional array) with axes *m/z*, *time*, and *sample*:
    a. m/z partitions with less than 5 time points in any sample are removed
    b. close m/z values are binned (tolerance chosen to match the experiment, e.g. 40 ppm and bin ends are rounded down to 4 decimal places)
    c. the tabular data is converted into a (m/z bin, scan cycle) matrix for each sample (scan file)
    d. the (m/z bin, scan cycle) of all samples matrices are stacked

#### Tensor Creation

To better align corresponding values across samples, and to reduce the size of the resulting tensor, binning is performed on both *m/z* and retention time axes. Spectra in a scan cycle will be offset forward on the time axis. Since each *m/z* vector ought to contain correlated values, we align the spectra belonging to the same cycle into the same matrix column (Fig. S1-3-1). This also helps reduce the size of the tensor. The *(m/z, cycle)* matrices for each sample are then stacked into a cube structure, taking care to align m/z and cycle indices and introduce missing values where no such index exists in any given sample. Missing values are imputed using Gaussian smoothing with a base value of zero.

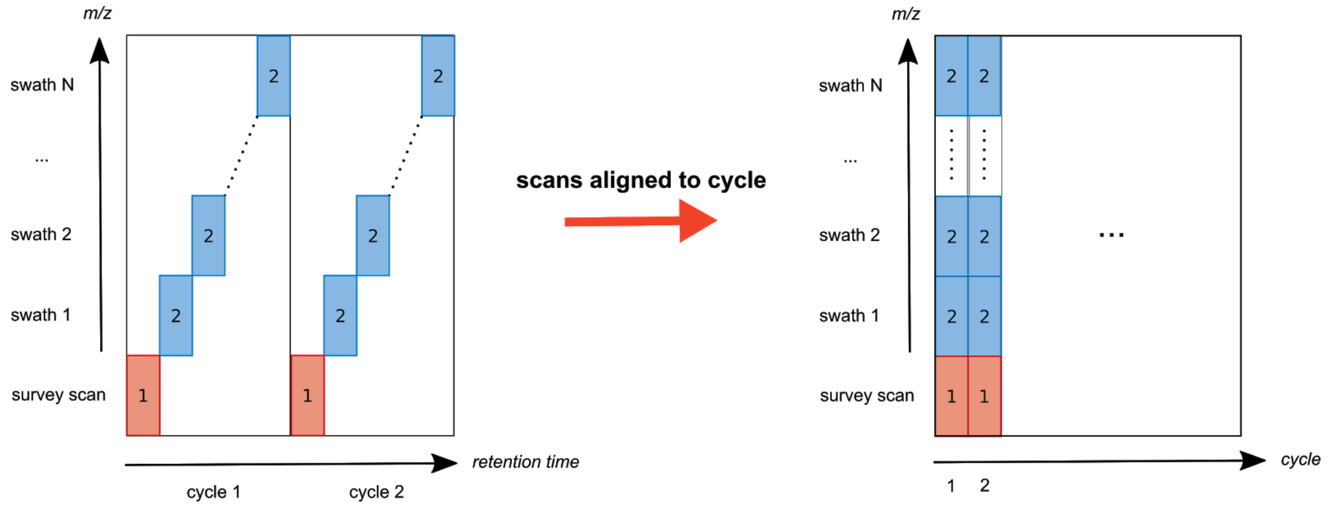

**Figure S1-3-1. Binning along the time axis when data tensors are created from SWATH-MS scan files.** Each MS 2 swath scan corresponding to a MS 1 survey scan has a delay of a few seconds, appearing as on the left after conversion of the scan files to matrices with binned time indices. All spectra that are part of the same cycle are aligned on a single column, and the time axis is replaced by the cycle index, and appear as on the right.

### Note S1-4. Processing of yeast lysate datasets

- Conversion to mzML was done with msconvert with peak picking enabled on both MS levels.
- The average elution peak width of 12 s (FWHM), used as a hint to the elution peak detection procedure, was estimated manually, by opening a scan in Spectronaut and determining the base width of iRT peptides.
- Conversion to CSV format was done with default pipeline parameters.
- The samples were sliced using a 1-minute retention time window. The width was considered appropriate given the full length of the chromatography and manual inspection of the number of peptides identified by Spectronaut in a window.
- Slice tensorization was performed with standard pipeline parameters.
- PARAFAC was run for a number of components F between 10 and 90 on the valid tensors.

### Note S1-5. Processing of the HYE110 (LFQbench) dataset

- Conversion to mzML was done with *msconvert* with peak picking enabled on both MS levels.
- The average elution peak width of 12.27 s (FWHM), used as a hint to the elution peak detection procedure, was estimated manually, by opening a scan in Spectronaut and determining the base width of iRT peptides.
- Conversion to CSV format was done with default pipeline parameters.
- The samples were sliced using a 5-minute retention time window. The width was considered appropriate given the full length of the chromatography and manual inspection of the number of peptides identified by Spectronaut in a window. 1664 slices were produced.
- Slice tensorization was performed with standard pipeline parameters. 1618 valid tensors were produced, with the rest discarded due to too little slice data.
- PARAFAC was run for a number of components F between 10 and 90 on the valid tensors. This yielded 147238 models.
- All model sample mode and time mode values were collected in tabular format files for downstream analysis
- The best models per slice were chosen using different criteria and the corresponding spectra were collected in a single pseudo-scan mzXML file for downstream analysis.

### Note S1-6. False positive underestimation assessment of DIA quantification

The expected number of false positive protein identifications, was computed as:

```
expected False Positive count =
FDR * mean(baseline count of protein IDs at that FDR across all tools) / median(pep:prot
ratio at that FDR)),
```

for each FDR threshold.

All mutant libraries were created using the same partially randomized FASTA file (each in silico Trypsin-digested fragment was shuffled). Skyline was run analogously with (Navarro et al. 2016), Spectronaut with default settings, except with 100% FDR threshold to allow selecting results at different levels. DIA-NN was run as below. Note that Spectronaut filters to only very low q-values, even if higher thresholds are set, hence the almost constant number of protein IDs across the different FDR thresholds.

#### Note S1-7. Assessment of information in unidentified components

To assess the correlation between PARAFAC components that were matched to peptides by either Crux or MS-GF+ and those which were not, a subsample of 1000 random unimodal components from both sets was chosen. The sample modes of these components were considered for the correlation. These consist of 30 points associated with the component contribution to each sample.

#### Note S1-8. Post-translational modifications search

We ran the MS-GF+ tool (Kim and Pevzner 2014) on PARADIAS output spectra resulting from the decomposition of the yeast replicates dataset to look for the following PTMs: acetylations, phosphorylations, succinylations, and GalNAc (core 1) glycosylations. The tool was configured for TOF instruments and allowed to account for a maximum of 384 modifications per peptide, resulting in much longer search time than the default value of 3. As a baseline comparison, we ran MS-GF+ on all DIA-Umpire output feature files separately, then collated the results. For this latter baseline run, there was almost no commonality of modified peptides found by MS-GF+ across the technical replicates, with only 8.2% appearing in at most 2 replicates Figure S1-6). Because of this, we considered median DIA-Umpire / MS-GF+ PTM counts across the replicates when comparing with PARADIAS / MS-GF+ results. Overall, we obtain roughly twice as many PTMs with MS-GF+ run on PARADIAS output.

### Note S1-9. The PARADIAS pipeline

The overall PARADIAS workflow consists of three main stages: preprocessing, decomposition, and output of spectra from best models. Additionally, to perform analyses on the output spectra, the pipeline provides a number of wrapper scripts to use interfaces with standard tools, as well as R and Jupyter notebooks to evaluate results. All stages may be run on either a workstation or a high-performance computing environment (the current implementation works with a Slurm-managed cluster). The decomposition relies on CUDA-capable GPU cards though it can be made to run on CPUs only, at a significant cost of speed per decomposition.

The pipeline is configured through a central YAML file that specifies the locations of the various files and directories, which third-party software to use, data-specific parameters (e.g. mass tolerance, size of the time window), as well as computational parameters (e.g. how much memory to use). Table S1-4 describes some parameters that are particularly important or whose meaning may not be clear from their name.

#### Preprocessing

In order for the scan files to be fed to the decomposition procedure, they must first be restructured into a tensor in (m/z, retention time, sample)-space. Note that we use “sample” and “scan file” interchangeably in this article since the latter is considered as a “sample” by the PARAFAC method. After conversion of scan files to a tabular format for easier processing, they are sliced according to swath and a predetermined retention time window. This takes advantage of the independence of isolation windows in terms of produced signals.

Each scan map in a slice is then converted to a (m/z, retention time) matrix, taking care to align entries on the m/z and RT axes. The m/z values are binned into partitions based on the set machine tolerance (e.g. 40 ppm) and differentiated by 4 decimal places. The RT dimension is binned into scan cycles and MS level 1 and level 2 spectra belonging to the same cycle are aligned into the same vector.

#### Decomposition

While in principle the full data tensor could be processed directly, the necessary memory is prohibitive (e.g. hundreds of gigabytes). Given the size of high-throughput SWATH LC—MS/MS data, it is necessary to divide the full tensor into independent partitions (“slices”). PARAFAC is run in parallel on all slice tensors, using the TensorLy library with PyTorch as a backend. By using GPUs and tensor processing frameworks, we take advantage of the massive parallelism and vectorized processing of graphics cards.

Each slice tensor is decomposed for all values of F in the preconfigured range. The Nvidia Multi-Processing Service (MPS) is leveraged to allow multiple parallel decompositions on the same GPU card. This is particularly efficient on newer cards with hardware MPS support (Volta models onwards). The current implementation was tested on the following Nvidia GPU card models: Tesla V100, Tesla K80, and Quadro GP100.

#### MS/MS file from best models

In this final processing step, the best models are selected from all that were generated, according to the criteria described in the Model selection section. The mass modes of these slice models, which hold deconvolved m/z spectra, are saved to an mzXML file. The mass mode (m/z spectrum) of a single component contains both MS level 1 and 2 intensities, corresponding to isolation window signals and their associated fragment signals, respectively. We are primarily concerned with MS level 2 spectra, and only record the highest intensity from the level 1 values. This highest MS 1 peak is set as the precursor of the MS 2 spectrum. Thus, the output file resembles DDA (shotgun) scans and DIA-Umpire output files, and may be processed with the same kind of tools that accept these as input. To remove residual noise from the decomposition procedure, the MS 2 intensities are baseline filtered by removing values that fall within the first 1% of that spectrum's intensity distribution. Each spectrum receives a unique index across the entire dataset, which may then be used to trace it back to its original slice.

Note that we do not record retention time information for these spectra, as this was not needed for the downstream analysis tools we used. This is however very easy to do by performing a cross product of the mass and time mode of each component. The resulting matrix is the LC-MS/MS map of that component and all spectra within it are copies of the mass mode spectrum with different scalings according to the time mode elution profile. These may be all saved then in the output mzXML file.

#### Analytics

Identification is performed using either the Crux toolkit, i.e. Comet for obtaining both target and decoy PSMs from PARADIAS output spectra, followed by Percolator confidence assignment, or MS-GF+, which also does confidence estimation. For quantification, we rely on a library quantification approach. We use the spectra output from PARADIAS to construct a library, following the protocol from (Schubert et al. 2015). We run the DIA-NN tool (Demichev et al. 2019) on the scan files using this library to get high-accuracy quantities. De novo sequencing is performed with Novor (Muth and Renard 2018) and DeepNovo (Tran et al. 2017).

#### Library Building Protocol

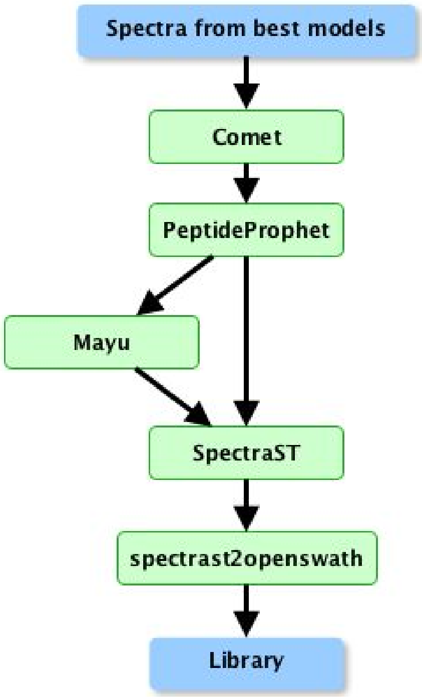

| Tool | Parameter | YAML variable or hard-coded value |
|---|---|---|
| **Comet** | peptide mass tolerance | 50 ppm |
| | database contents | concatenated target and decoy sequences |
| **Mayu** | FDR (`-G`) | `quant_lib_mayu_fdr` |
| | number of analysis steps (`-H`) | 51 |
| | filtering (`-P`) | filter data with PSM FDR `quant_lib_mayu_fdr` (e.g. mFDR=0.1:t) |
| **SpectraST** | fragmentation type (`-cI`) | CID-QTOF |
| | minimum probability to include (`-cP`) | 0.05 |
| | remove spectra of decoys (`-c_RDY`) | Yes |
| **spectrast2 openswath (spectrast2tsv)** | fragment m/z range (`-l`) | `lower_mz_frag`, `upper_mz_frag` |
| | ion types considered (`-s`) | b,y |
| | charges considered (`-x`) | 2, 3 |
| | minimum number of reported ions per peptide/z (`-o`) | 4 |
| | maximum number of reported ions per peptide/z (`-n`) | 12 |
| | maximum error allowed at the annotation of a fragment ion (`-p`) | `quant_library_spectrast_max_frag_annot_err` |
| | removed duplicate masses from labeling (`-d`) | Yes |
| | used theoretical mass (`-e`) | Yes |

### Note S1-10. Software Versions

Below is a list of the software versions used in this study.
For an exhaustive specification of all dependencies, see the PARADIAS GitHub repository.

- ProteoWizard 3.0.10462 (msconvert)
- Crux version 3.2-0bf523f2
- MS-GF+ v2019.07.03
- DIA-Umpire 2.0
- TPP v5.2.0 Flammagenitus, Build 201904252338-7913
- Mayu 1.08
- msproteomicstools (SpectraST) 0.11
- DIA-NN 1.7.4
- Skyline 20.1.0.31
- Spectronaut 13.10.191212.43655
- DeepNovo 0.0.1
- Novor 1.06.0634
- Python (3.6.7) packages
  - TensorLy 0.4.2
  - PyTorch 1.1.0
  - SciPy 1.2.1
  - NumPy 1.16.4
  - AstroPy 3.0.5
- R (3.6.2) packages
  - tidyverse 1.3.0
  - data.table 1.12.8
- Snakemake 5.7.4

Harshman, Richard A. 1970. *Foundations of the PARAFAC Procedure: Models and Conditions for an “Explanatory” Multi-Modal Factor Analysis*. University of California at Los Angeles.

Håstad, Johan. 1989. “Tensor Rank Is NP-Complete.” In *Automata, Languages and Programming*, 451–60. Springer Berlin Heidelberg.

Johnsen, L. G., J. M. Amigo, and T. Skov. 2014. “Automated Resolution of Overlapping Peaks in Chromatographic Data.” *Journal of*. https://onlinelibrary.wiley.com/doi/abs/10.1002/cem.2575.

Kim, Sangtae, and Pavel A. Pevzner. 2014. “MS-GF+ Makes Progress towards a Universal Database Search Tool for Proteomics.” *Nature Communications* 5 (October): 5277.

Kolda, Tamara G., and Brett W. Bader. 2009. “Tensor Decompositions and Applications.” *SIAM Review* 51 (3): 455–500.

Liu, Kefei, João Paulo C. L. da Costa, Hing Cheung So, Lei Huang, and Jieping Ye. 2016. “Detection of Number of Components in CANDECOMP/PARAFAC Models via Minimum Description Length.” *Digital Signal Processing* 51 (April): 110–23.

Ma, Bin. 2015. “Novor: Real-Time Peptide de Novo Sequencing Software.” *Journal of the American Society for Mass Spectrometry* 26 (11): 1885–94.

Muth, Thilo, and Bernhard Y. Renard. 2018. “Evaluating de Novo Sequencing in Proteomics: Already an Accurate Alternative to Database-Driven Peptide Identification?” *Briefings in Bioinformatics* 19 (5): 954–70.

Navarro, Pedro, Jörg Kuharev, Ludovic C. Gillet, Oliver M. Bernhardt, Brendan MacLean, Hannes L. Röst, Stephen A. Tate, et al. 2016. “A Multicenter Study Benchmarks Software Tools for Label-Free Proteome Quantification.” *Nature Biotechnology* 34 (11): 1130–36.

Schubert, Olga T., Ludovic C. Gillet, Ben C. Collins, Pedro Navarro, George Rosenberger, Witold E. Wolski, Henry Lam, et al. 2015. “Building High-Quality Assay Libraries for Targeted Analysis of SWATH MS Data.” *Nature Protocols* 10 (3): 426–41.

Skov, Thomas, and Rasmus Bro. 2008. “Solving Fundamental Problems in Chromatographic Analysis.” *Analytical and Bioanalytical Chemistry* 390 (1): 281–85.

Smilde, Age, Rasmus Bro, and Paul Geladi. 2005. *Multi-Way Analysis: Applications in the Chemical Sciences*. John Wiley & Sons.

Tran, Ngoc Hieu, Xianglilan Zhang, Lei Xin, Baozhen Shan, and Ming Li. 2017. “De Novo Peptide Sequencing by Deep Learning.” *Proceedings of the National Academy of Sciences of the United States of America* 114 (31): 8247–52.

Vowinckel, Jakob, Aleksej Zelezniak, Roland Bruderer, Michael Mülleder, Lukas Reiter, and Markus Ralser. 2018. “Cost-Effective Generation of Precise Label-Free Quantitative Proteomes in High-Throughput by microLC and Data-Independent Acquisition.” *Scientific Reports* 8 (1): 4346.
